# Melt Electrowritten Scaffold-Reinforced Affibody-Conjugated Hydrogels for Controlled Bone Morphogenetic Protein-2 Delivery

**DOI:** 10.1101/2025.10.14.682494

**Authors:** Jonathan Dorogin, Yan C. Pacheco, Patrick C. Hall, Andy J Huang, Payton M. Jefferis, Kaitlyn A. Link, Morrhyssey A. Benz, Paul D. Dalton, Nick J. Willett, Marian H. Hettiaratchi

## Abstract

Bone morphogenetic protein-2 (BMP-2) is clinically used to promote bone regeneration but suffers from uncontrolled release when delivered from collagen sponges, necessitating high doses that can cause adverse effects. Hydrogels offer tunable protein release but are limited by weak mechanics and poor stability during storage and handling. Here, we introduce a two-part protein delivery platform that integrates mechanical reinforcement with affinity-controlled protein release. We developed a melt electrowritten (MEW) scaffold–reinforced, affibody-conjugated polyethylene glycol maleimide (PEG-mal) hydrogel for affinity-controlled BMP-2 delivery. MEW scaffolds improved hydrogel handling, compressive resistance, and stability during lyophilization and rehydration, without altering bulk stiffness. Engineered BMP-2-specific affibodies provided affinity-based control over BMP-2 release. This ability to control BMP-2 release was preserved after lyophilization and rehydration of the hydrogels. *In vivo*, affibody conjugation of high-affinity affibodies to the hydrogels significantly enhanced BMP-2 retention in subcutaneous implants, while MEW reinforcement significantly increased bone volume and defect bridging in rat femoral bone defects. This affibody-conjugated, MEW scaffold-reinforced hydrogel system effectively integrates mechanical reinforcement with tunable protein-material affinity interactions, advancing hydrogel-based delivery strategies for BMP-2 and other protein therapeutics in musculoskeletal repair.

## 1. Introduction

Spinal degeneration is an increasingly common medical diagnosis, resulting in over 450,000 annual spinal fusion surgeries in the United States alone.^1^ Due to their complexity and prevalence, spinal fusion surgeries account for the greatest aggregate cost for operating room procedures, amounting to $14.1 billion in 2018 in the United States.^2^ Although spinal fusion procedures have a relatively high success rate, up to 70% of patients have reported post-surgical complications,^3^ with 24% requiring revision surgeries that significantly impact patients’ quality of life.^4^ While the cause of spinal fusion failure is typically multi-factorial, common themes include the failure of fixation materials,^5^ excessive inflammation,^3,6^ and inadequate bone healing.^6,7^ Therefore, exogenous supplementation of one or more proteins to restore the events of the bone healing cascade is a promising therapeutic approach for stimulating bone repair.^8^ Bone morphogenetic protein-2 (BMP-2) is an osteogenic protein often delivered in supraphysiological concentrations for the repair of large bone defects, especially in the spinal column.^3,9,10^ In the clinic, BMP-2 is commonly delivered from an absorbable collagen sponge encased in a plastic or metal spinal cage.^11^ However, BMP-2 interacts with collagen through weak electrostatic interactions, which can result in poorly controlled, rapid protein release from collagen-based biomaterials. This uncontrolled BMP-2 release has been associated with side effects such as ectopic and heterotopic ossification,^12^ a heightened inflammatory response, and swelling,^13^ necessitating the development of biomaterials that can better spatially and temporally localize protein delivery.^14,15^ Furthermore, biomaterials commonly used for pre-clinical BMP-2 delivery include from collagen, alginate, or polyethylene glycol (PEG) hydrogels that may not be able to withstand the natural mechanical loading environment of bone tissue and therefore must be delivered in a load-protected environment.^16^ This combination of factors poses a unique engineering challenge, as the biomaterial delivery vehicle must be sufficiently soft and porous to allow cell penetration and remodeling (<1 kPa)^17–20^ and simultaneously mechanically stable to resist premature deformation and degradation within the bone injury environment.^21^ Because protein delivery from biomaterials is often mediated by material porosity and mechanical properties, decoupling protein delivery rate from biomaterial mechanical integrity poses an additional challenge.^19,22^ For clinical translation, the engineered solution should also be easy to use by clinicians^23^ and amenable to long-term shelf stability.^24^ Developing a protein delivery strategy that effectively overcomes all of these limitations would be a significant innovation for the treatment of large bone defects, potentially enabling the expanded use of clinical BMP-2 treatment to other types of bone injuries.

Protein delivery vehicles that regulate protein release via electrostatic and hydrophobic affinity interactions between the protein and the material present an attractive alternative to protein release mediated by diffusion and material degradation. In these materials, the affinity strength of the protein-material interactions can be tuned to adjust protein release rate without drastically changing the mechanical properties of the delivery vehicle.^25^ Affinity-based biomaterials fabricated from natural polymers that participate in affinity interactions with proteins, such as heparin, have been shown to improve the local retention of therapeutic proteins such as BMP-2.^14,26–29^ More recently, synthetic binding proteins have been engineered with specific affinities for therapeutic proteins and incorporated into biomaterials at varying ratios to control protein release.^30–33^ We have previously engineered binding proteins called affibodies, a class of small, thermally stable protein binders derived from the z-domain of staphylococcal protein A,^34–38^ with high specificity and a range of affinities for several therapeutic proteins, including vascular endothelial growth factor (VEGF), platelet-derived growth factor (PDGF), fibroblast growth factor-2 (FGF-2), and BMP-2.^39–41^ We demonstrated that BMP-2 release from a PEG-based hydrogel could be tuned by conjugating affibodies with different affinities for BMP-2 to the biomaterial. Based on these *in vitro* results, we hypothesized that BMP-2-specific affibody-conjugated PEG hydrogels could be similarly used to locally control BMP-2 release *in vivo* and stimulate bone regeneration.

The challenge of mechanical stability of a hydrogel for protein delivery could be addressed by increasing the bulk hydrogel stiffness using higher polymer or crosslinker concentrations or reinforcing the hydrogel with other materials.^42^ However, increasing the bulk stiffness of the hydrogel could unintentionally impact many aspects of the material’s interactions with its surrounding environment, including cell-material and protein-material interactions, biomolecule diffusion, material degradation, and tissue remodeling.^17,19^ Alternatively, mechanical reinforcement with a secondary material could be applied without altering the hydrogel formulation itself, retaining many of its desired characteristics.^43^ This approach is similar to the clinical use of BMP-2 for spinal fusion, in which a soft absorbable collagen sponge is caged in PEEK^44^ or titanium^45^ to provide additional mechanical stability and prevent implant migration.^3^ However, these traditional devices for containing hydrogels and sponges for therapeutic delivery *in vivo* are stiff and bulky, interfering with tissue integration and causing mechanical mismatches with the surrounding environment. For BMP-2 delivery, forcibly manipulating the collagen sponge into the cage can squeeze out the BMP-2 solution, introducing variability between procedures and surgeons.

Building upon this concept, recent advances in additive manufacturing can be leveraged to integrate reinforcements into a hydrogel delivery vehicle instead of encasing it entirely, maintaining desired interactions between the hydrogel and its surrounding environment.^43^ Melt electrowriting (MEW) is an emerging, high-precision 3D printing technique that can generate highly porous scaffolds composed of ordered microfibers using clinically relevant thermoplastics such a poly ε-caprolactone (PCL), poly(lactic acid), and polypropylene.^46–49^ MEW relies on electrohydrodynamic forces to stabilize and precisely deposit thin fibers of molten polymer into defined shapes using tool paths similar to traditional extrusion-based 3D printing. Due to the applied electrohydrodynamic forces, MEW fibers are typically on the scale of 1-50 µm, MEW scaffolds retain porosity of 80-95% between fibers, and varying scaffold designs lead to broad ranges of mechanical properties.^50–52^ For instance, the combination of MEW scaffolds with methacrylated gelatin hydrogels resulted in a 54-fold increase in the stiffness of the composite construct.^43^ Unlike typical extrusion 3D printing that produces thicker, less flexible fibers, the microfibers and high scaffold porosity of MEW allow for increased flexibility, improving the ability to handle and surgically implant low modulus hydrogels without damaging the materials or surrounding tissues. Furthermore, hydrogels can be crosslinked directly within MEW scaffolds to avoid extra manipulation that may damage the materials before surgical implantation. Thus, we hypothesized that MEW reinforcement of affibody-conjugated hydrogels would increase hydrogel mechanical stability such that the hydrogels could be easily handled during surgery and remain localized within the implantation site over the period of time required for BMP-2 delivery, thereby improving BMP-2 retention without substantially affecting the mechanical microenvironment or impeding cell/tissue infiltration during healing.

A critical design criterion for clinical translation of a protein delivery vehicle is its long-term shelf stability, which can drastically reduce costs associated with maintaining a temperature-controlled supply chain and improve ease of use in the clinic.^53,54^ Drying materials through lyophilization is a common approach for increasing a biological product’s shelf stability.^53,55^ For example, the clinical BMP-2 delivery device consists of a lyophilized collagen sponge that is rehydrated with a highly concentrated solution of BMP-2 prior to surgical implantation. However, lyophilization can damage the crosslinked polymer network of hydrogels, resulting in material degradation, altered mechanical properties, and unpredictable protein release,^55,56^ or induce protein unfolding, denaturation, and aggregation, reducing its therapeutic efficacy.^24^ Therefore, determining the effect of lyophilization on protein delivery from affibody-conjugated hydrogels may provide valuable insights into their future potential for shelf stability and clinical translation. We hypothesized that reinforcement with the MEW scaffold would help maintain the structural integrity of the hydrogel during the lyophilization and rehydration processes. We further hypothesized that affinity interactions between BMP-2 and the affibody-conjugated hydrogels would not be impacted by lyophilization and rehydration of the hydrogels.

In this work, we engineered an affinity-based delivery system for BMP-2 with improved mechanical stability for surgical implantation and the potential for lyophilization for long-term shelf stability. We reinforced affibody-conjugated PEG hydrogels with MEW scaffolds to increase their mechanical stability and integrity without altering the hydrogel formulation and subjected the scaffold-reinforced hydrogels to mechanical testing to identify the impact of reinforcement on the hydrogel’s physical properties. We integrated affibodies into the hydrogels to control the release of BMP-2 and investigated the effect of the MEW scaffold on protein release. The reinforced hydrogels underwent lyophilization and subsequent rehydration followed by additional mechanical testing and protein release experiments to determine the impact of lyophilization and rehydration on the performance of the delivery system. The effect of affibody incorporation in the hydrogels on BMP-2 retention *in vivo* was also evaluated using fluorescence imaging. Finally, PEG hydrogels without reinforcement or affibodies, with reinforcement, and with reinforcement and affibodies were loaded with BMP-2 and implanted in a critically sized femoral bone defect in rats to evaluate their efficacy in stimulating bone repair over 12 weeks. This rodent bone injury model is a well-characterized, critically sized defect in a load-bearing bone that enables longitudinal assessment of bone formation via x-ray and micro-computed tomography.^14,57,58^ It provides an excellent initial testbed to investigate the use of MEW scaffold reinforcement and BMP-2-specific affibodies for improving BMP-2 delivery for bone regeneration prior to investigation in larger animal models of spinal fusion. Together, these studies demonstrate the development and use of a novel delivery system for affinity-controlled delivery of BMP-2 for clinically relevant bone repair.

## 2. Materials and Methods

### 2.1. Affibody synthesis

High-affinity (dissociation constant, K_D_ = 10.7 nM) and low-affinity (K_D_ = 34.8 nM) BMP-2-specific affibodies were either expressed recombinantly in *E. coli* and purified using immobilized metal affinity chromatography followed by size exclusion chromatography^39^ or synthesized via solid-phase peptide synthesis and purified using high performance liquid chromatography.^59^

For recombinant affibody expression, BMP-2-specific affibody sequences modified with a C-terminal hexahistidine tag and cysteine were ligated into the pET28b+ expression vector for isopropyl β-D-1-thiogalactopyranoside (IPTG)-inducible protein expression. The plasmids were transformed into chemically competent BL21(DE3) *E. coli* (New England Biolabs, Ipswich, MA, United States). Transformed *E. coli* were grown in Luria-Bertani broth (Fisher BioReagents, Waltham, MA, United States) supplemented with 50 mg/L of kanamycin sulfate (Millipore Sigma, Burlington, MA, United States) overnight followed by growth in Terrific Broth (Research Products International, Mount Prospect, IL, United States), supplemented with 50 mg/L of kanamycin sulfate and 400 µL of Antifoam 204 (Millipore Sigma, Burlington, MA, United States) at 37 °C in a water bath bioreactor (Epiphyte3, Toronto, ON, Canada) with air sparging until an optical density at 600 nm of approximately 1.2 was reached. The *E. coli* were then induced to produce affibodies by adding IPTG (Gold Biotechnology) to a final concentration of 0.5 mM and incubating the culture at 18 °C for 20 hours. The cells were lysed by sonication in 50 mM tris buffer (Fisher Scientific) supplemented with 500 mM NaCl (Fisher Scientific), 5 mM imidazole (Fisher Scientific), and 75 mg of tris(2-carboxyethyl)phosphine (TCEP) (Gold Biotechnology). The lysate was centrifuged, and the supernatant was collected and mixed with cobalt-nitrilotriacetic acid (Co-NTA) (Gold Biotechnology) beads for 60 minutes to form a chelation bond between the cobalt and the hexahistidine tag of the affibodies. The solution was transferred to an empty glass chromatography column (BioRad, Hercules, CA, United States) and washed with 50 mM tris buffer containing 500 mM NaCl (Fisher Scientific) and 30 mM imidazole (Fisher Scientific) to remove weakly bound proteins. The purified affibodies were then eluted from the column with 50 mM tris buffer containing 500 mM NaCl and 250 mM imidazole. Affibodies were buffer-exchanged into phosphate-buffered saline (PBS) (Fisher Scientific) and further purified by size exclusion chromatography (BioRad NGC with Enrich SEC 70, 10/300 mm column; BioRad). Affibodies were stored at -20 °C until use.

Solid-phase peptide synthesis of affibodies was performed using a Liberty Blue 2.0 microwave peptide synthesizer (CEM Corporation, Matthews, NC, United States). The high- and low-affinity BMP-2-specific affibody sequences were modified with a penultimate cysteine and a C-terminal glycine. Solutions of 0.2 M of each amino acid (CEM Corporation), Fmoc-Gly-Wang resin (CEM Corporation) (0.602 mmol/g loading), 10% (v/v) pyrrolidine (Millipore Sigma), 1 M N,N’-diisopropylcarbodiimide (DIC) (Oakwood Chemical), and 1 M Oxyma (CEM Corporation) were prepared in N,N-dimethylformamide (DMF) (Fisher Scientific) and loaded onto the peptide synthesizer. Synthesized crude affibodies were cleaved from the resin by mixing with a cleavage cocktail (0.5 mL H_2_O, 0.5 mL triisopropyl silane (Oakwood Chemical, Estill, SC, United States), 0.5 mL 2,2’-(ethylenedioxy)diethanethiol (Tokyo Chemical Industry America, Portland, OR, United States), 0.5 mL thioanisole (Oakwood Chemical), 18 mL trifluoroacetic acid (TFA) (Oakwood Chemical)) for 40 minutes at 42 °C. The cleaved solution was vacuum filtered, and the filtrate was purified via precipitation and three rounds of centrifugation (1200 RCF, 5 minutes) using cold diethyl ether (Fisher Scientific). The precipitate was dried overnight in a desiccator and purified using a Prodigy high-performance liquid chromatography (HPLC) system (CEM Corporation) by running a 30-75% gradient of acetonitrile in H_2_O with 0.1% (v/v) TFA. Affibodies were lyophilized and stored at -20 °C until use.

The size and purity the recombinant and synthetic affibodies were confirmed using matrix-assisted laser desorption/ionization time of flight (MALDI-TOF) mass spectrometry (Bruker Smart LS; Bruker, Billerica, MA, United States), and the secondary structures of the affibodies were confirmed using circular dichroism (JASCO J-815; JASCO Corporation, Tokyo, Japan), as previously described.^59^

### 2.2. Poly ε-caprolactone scaffold fabrication

Tubular MEW PCL scaffolds were printed with a diamond pore design using a custom-built, high-precision MEW system based on the Aerotech A3200 platform (Aerotech, Inc., Pittsburgh, PA, United States) with a cylindrical collector.^51,52^ The designed scaffold had a programmed diameter of 5 mm, a pore area of 0.5 mm^2^, and a layer count of 10 fibers. The tubular MEW scaffolds had lengths of 5, 7.5, or 10 mm and internal volumes of 100, 150, or 200 μL, respectively. Within MEW, the material deposition lags the nozzle and is different for different parameters, including printer type. The toolpath movement was adjusted to result in MEW scaffolds with the sizes mentioned above. MATLAB (MathWorks) was used to write a scaffold toolpath code to translate these input parameters into the AeroBasic programming language.^51,65^

PCL (Corbion PURASORB PC12, Lot #2007001491; Corbion, Amsterdam, Holland) was loaded into 3 mL glass syringes (Fortuna Optima, Poulten & Graf GmbH, Wertheim, Germany), which were capped with 22-gauge nozzles (Vita Needle Company, Needham, MA, United States) and heated to 80 °C overnight to allow for the PCL to melt to the bottom of the nozzle and release trapped air bubbles. Then, the loaded syringe was placed into a heating jacket and fitted with a pressure connector. An input pressure of 75 kPa, hot-end temperature of 80 °C, collector translation speed of 4.5 mm/s, collector distance of 3 mm, applied voltage of 4.5 kV, and mandril diameter of 5 mm were used to print the scaffolds.

The 5 mm long scaffold was used to contain a 100 µL hydrogel for *in vitro* experiments, the 7.5 mm long scaffold was used to contain a 150 µL hydrogel for subcutaneous implants, and the 10 mm long scaffold was used to contain a 150 µL hydrogel for placement in the femoral bone defect.

### 2.3. Physical characterization of tubular MEW scaffolds

Optical and scanning electron microscopy (SEM) were used to visualize the physical structure of the MEW scaffolds. A Keyence VHX-7000 microscope was used for optical imaging of the scaffolds to analyze the structure and fiber diameter of the scaffolds. For SEM, the MEW scaffolds were first sputter-coated using an Au-Pd alloy (Cressington) prior to imaging on a ThermoFisher Apreo 2 SEM using 3.2 nA, 5.00 kV, and 39-1000x magnification.

### 2.4. Synthesis of affibody-conjugated PEG-mal hydrogels

Affibody-conjugated polyethylene glycol-maleimide (PEG-mal) hydrogels were synthesized at volumes of either 100 or 150 µL (5%, w/v) as previously described.^39,66^ Briefly, 20 kDa, 4-arm PEG-mal (Laysan Bio, Inc., Arab, AL, United States), high- or low-affinity BMP-2- specific affibodies, and/or cell-adhesive RGD peptides (CGRGDSG, prepared and purified in-house using the CEM Liberty Blue 2.0 and CEM Prodigy HPLC described for affibody synthesis) were each dissolved in PBS pH 6.9 and mixed for 30 minutes at room temperature to form PEG, PEG-affibody, PEG-RGD, or PEG-affibody-RGD intermediates. The intermediate solutions were crosslinked in 2 mL centrifuge tubes with dithiothreitol (DTT) (Gold Biotechnology) dissolved in PBS pH 6.9 for 30 minutes at room temperature. Each hydrogel contained 500 times molar excess of affibody compared to the amount of BMP-2 loaded for each application and 1 mM RGD peptide for *in vivo* implantation. Unreacted DTT was removed from the hydrogels prior to loading with BMP-2 by washing the hydrogels 3 times with PBS.

### 2.5. Calculation of hydrogel mesh size

Hydrogel mesh sizes were calculated using the Equilibrium Swelling Theory.^22,67–69^ 100 µL 5% (w/v) PEG-mal hydrogels were synthesized without affibodies, with 2 nmol of high-affinity BMP-2-specific affibodies, and/or with 10 µmol of RGD peptides and crosslinked with 45-50 µmol of DTT based on the formulation. The initial masses of the hydrogels were recorded immediately after crosslinking (i.e., the relaxed state). The hydrogels were then swollen overnight in 1 mL of PBS at 4 °C. The PBS solution was removed, and the hydrogels were weighed again in the swollen state. The hydrogels were then frozen at -80 °C overnight and lyophilized at -104 °C and 50 mTorr for 12 hours using a SP Scientific ZG benchtop lyophilizer, and their dry masses were measured. The hydrogels were also rehydrated with 100 µL of PBS, and the process of swelling and lyophilizing was repeated to determine if hydrogel mesh sizes were affected by lyophilization. **Equation 1** and **Equation 2** were used to determine the approximate mesh size of the hydrogels, where *M_n_* is the number-average degree of polymerization of the 4-arm PEG-mal (20,000), υ. is the specific volume of bulk PEG-mal (0.888 cm^3^/g, calculated as the reciprocal of the density of 4-arm PEG-mal (1.125 g/cm^3^)), *V_1_* is the molar volume of water (18 mL/mol), υ*_2,s_* is the equilibrium swollen polymer volume fraction, υ*_2,r_* is the relaxed polymer volume fraction, χ is the Flory-Huggins interaction for PEG in water (0.426),^67^ *C_n_* is the characteristic ratio of PEG (3.96),^70^ *M_r_* is the molecular weight of PEG repeats (44 g/mol), and *l* is the bond length of the PEG backbone (0.146 nm).^67^

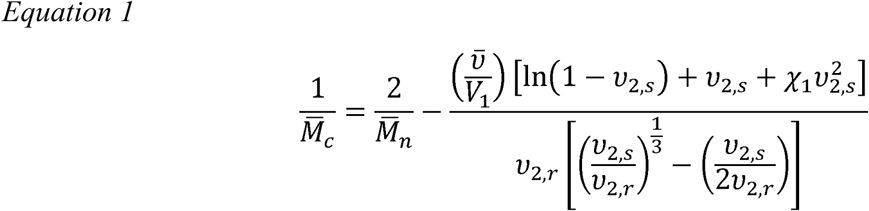

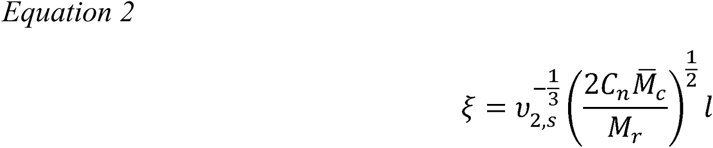

### 2.6. Fabrication of scaffold-reinforced hydrogels

To form PEG-mal hydrogels within the MEW scaffolds, the scaffold was held horizontally, and the PEG intermediate solution was pipetted into the interior of the tubular scaffold (**Supplemental Video S1**). The DTT solution was then added dropwise to the interior of the scaffold to crosslink the hydrogel, after which the MEW scaffold was rotated such that the solutions would uniformly fill the interior of the tube. The scaffold was suspended for at least 10 minutes to allow the DTT to fully crosslink the available maleimides. The scaffold-reinforced hydrogels were then transferred to 2 mL centrifuge tubes and submerged in 1 mL of PBS. The hydrogels were washed 3 times with fresh PBS to remove unreacted DTT.

### 2.7. Swelling and stability of hydrogels

Hydrogels with and without MEW scaffolds were swollen overnight in 1 mL of 0.1% bovine serum albumin (BSA; Fisher Scientific) in PBS in 2 mL microcentrifuge tubes and incubated at 37 °C for 14 days. Images were taken before swelling and at 7 and 14 days. Hydrogels were weighed periodically over 14 days, and the mass of the MEW scaffold was subtracted.

### 2.8. Lyophilization of hydrogels

Scaffold-reinforced hydrogels with and without affibodies were lyophilized to investigate the effect of lyophilization on hydrogel mechanical properties and protein release. Hydrogels were transferred to a dry petri dish and flash-frozen using liquid nitrogen. The frozen hydrogels were lyophilized at -104 °C and 50 mTorr for 12 hours using a SP Scientific ZG benchtop lyophilizer. After lyophilization, the hydrogels were stored at 4 °C for 72 hours until rehydration for assessment of mechanical and rheological properties and protein release kinetics. Hydrogels were rehydrated with 130 µL of PBS.

### 2.9. Mechanical and rheological characterization of reinforced and lyophilized hydrogels

Radial compressive testing was performed on non-reinforced hydrogels, MEW scaffolds, and scaffold-reinforced hydrogels using a DHR-2 hybrid rheometer (TA Instruments, New Castle, DE, United States). 100 µL 5% (w/v) PEG-mal hydrogels were cast in 8 mm diameter silicone molds to form discs, which were used to investigate the baseline mechanical properties of the hydrogels. Materials were compressed radially by 98% of their original width at a rate of 10 µm/s. Stiffness was calculated as the slope of the stress vs. strain curve to 10% compression. Compressive testing was also performed on scaffold-reinforced hydrogels that had undergone a cycle of lyophilization and rehydration. Lyophilized hydrogels were rehydrated in PBS 30 minutes prior to testing.

### 2.10. BMP-2 release from hydrogels

To assess the effects of the MEW scaffold on protein release, affibody-conjugated hydrogels were fabricated with and without MEW scaffold reinforcement, washed to remove excess DTT, and loaded with 100 ng (4×10^-12^ mol) of BMP-2 (R&D Biotechne, Minneapolis, MN, United States). Hydrogels were synthesized without affibodies, with 2 nmol of low-affinity BMP-2-specific affibodies, or with 2 nmol of high-affinity BMP-2-specific affibodies to achieve 500 molar excess of affibody to BMP-2, which we have demonstrated reduces the rate and total amount of protein released from PEG-mal hydrogels.^39^ The hydrogels were crosslinked with 50 µmol of DTT using the remaining available maleimides. 5 µg/mL BMP-2 in PBS were added to the hydrogels for 30 minutes to mimic clinical protein loading protocols.^71^ 200 µL of 0.1% (w/v) BSA in PBS were then added each hydrogel and immediately removed to quantify the amount of BMP-2 loaded (i.e., BMP-2 encapsulation efficiency). 1 mL of 10% (v/v) fetal bovine serum (FBS; Gibco, Thermo Fisher) in PBS was added to each hydrogel, and 200 µL of FBS solution were immediately removed to quantify initial protein release. 200 µL of fresh FBS solution was replaced, and 200 µL samples were similarly taken for 7 days. For comparison to a clinically relevant biomaterial, cylindrical collagen sponges were prepared from a collagen sponge sheet (Medtronic Infuse™ bone graft kit) using an 8 mm biopsy punch (Integra™ Miltex™), loaded with 100 ng of BMP-2, and allowed to release BMP-2 into 10% (v/v) FBS solution over 7 days with periodic sampling. We chose to evaluate BMP-2 release for 7 days based on our previous results,^39^ which indicated a plateau in BMP-2 release at time points taken after 7 days. BMP-2 encapsulation and release were quantified using a BMP-2-specific enzyme linked immunosorbent assay (ELISA) (R&D Biotechne).

In addition to cumulative protein release over time, the protein release profiles were fit to the Korsmeyer-Peppas model (**Equation 3**), where M_t_/M_∞_ is the fraction of BMP-2 released at time t (in seconds), K is the effective diffusivity, and n is the diffusion exponent that indicates the mechanism of transport.^72,73^ Data that were M_t_/M_∞_≤ 0.6 were fitted, and the effective diffusivity (K) of BMP-2 from each hydrogel formulation was identified as the slope of the curve.^72,73^

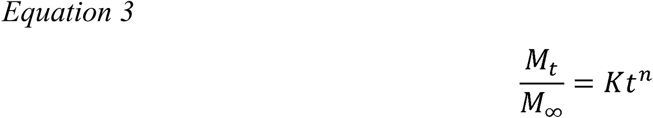

To assess the effects of lyophilization on protein release, MEW-reinforced hydrogels were fabricated as described without affibodies, with 2 nmol of low-affinity BMP-2 affibodies, or with 2 nmol of high-affinity BMP-2 affibodies. A control group was stored in PBS at 4 °C for 48 hours, and a treatment group was flash frozen, lyophilized, and stored dry for 48 hours. Hydrogels were then transferred into dry 2 mL microcentrifuge tubes and loaded with 100 ng of BMP-2 for 30 minutes. For control hydrogels, a 20 µL of 5 µg/mL BMP-2 in PBS was added, and for the lyophilized groups, the hydrogels were fully rehydrated in a 130 µL solution of 0.769 µg/mL BMP-2 in PBS. BMP-2 encapsulation and release were similarly assessed as described above.

### 2.11. Subcutaneous implantation of scaffold-reinforced hydrogels

MEW scaffold-reinforced hydrogels were implanted in the dorsal subcutaneous space of 5-6 week old female Sprague Dawley rats (Charles River Laboratories, Wilmington, MA, United States) to investigate local protein retention *in vivo*. All animal procedures were approved by and conducted in accordance with the University of Oregon’s Institutional Animal Care and Use Committee (IACUC). All reagents were sterile-filtered through a 0.22 µm filter and handled in a biosafety cabinet using aseptic techniques. BMP-2 was fluorescently labeled with IRDye 800CW NHS Ester (LI-COR Biosciences, Lincoln, NE, United States) according to the manufacturer’s protocols. The fluorescent degree of labeling was determined to be an average of 3.86 fluorescent molecules per protein. 150 µL 5% (w/v) affibody-conjugated PEG-mal hydrogels were prepared in 7.5 mm long MEW scaffolds. The hydrogels were functionalized with 1 mM RGD to promote cell adhesion and infiltration^74,75^ and 38 nmol of high- or low-affinity BMP-2-specific affibodies. The hydrogels were crosslinked with 43.1 µmol or 45 µmol of DTT based on the remaining available maleimides and loaded with 2 µg of fluorescent BMP-2. 20 µL of 0.1 mg/mL BMP-2 were added on top of the hydrogels and allowed to soak into the hydrogel in the dark for 3 hours.

The fur on the dorsal aspect of the rats was shaved, and the surgical site was aseptically prepared using 70% isopropyl alcohol (VetOne) and 4% chlorhexidine solution (VetOne). The rats were anesthetized using inhalant isoflurane anesthesia (VetOne). Two 1.5 cm incisions were made along the dorsum of each rat, one midline between the shoulder blades and the other just cranial to the pelvis. Blunt dissection was used to create two subcutaneous pockets per incision.^76^ Hydrogels were implanted in the subcutaneous pockets, and the incisions were closed with wound clips. The locations of the hydrogels from each treatment group were randomized.

After 21 days, the rats were euthanized via carbon dioxide asphyxiation, following the recommendations of the American Veterinary Medical Association (AVMA) Guidelines for the Euthanasia of Animals. The hydrogels were explanted from the subcutaneous space and transferred to a 24 well plate. Fluorescent images of the hydrogels were captured using a Spectrum In Vivo Imaging System (IVIS) (Perkin Elmer, Inc., Waltham, MA, United States) with 745 nm excitation and 800 nm emission wavelengths and exposure time of 2 seconds. To quantify fluorescent signal, regions of interest (ROIs) were drawn around each hydrogel, total radiant efficiency was measured, and background signal was subtracted.

### 2.12. Femoral bone defect surgery

PEG-mal hydrogels without scaffold reinforcement or affibodies, with scaffold reinforcement, and with scaffold reinforcement and high-affinity BMP-2 affibodies were implanted in critically sized femoral bone defects in rats to investigate the effect of scaffold reinforcement and affibodies on BMP-2-induced bone repair. 150 µL 5% (w/v) PEG-mal hydrogels were synthesized with or without scaffold reinforcement, with 1 mM RGD peptide, and with or without 96 nmol of high-affinity BMP-2 affibodies. 10 mm long MEW scaffolds were used such that the unfilled ends of the scaffold encircled the bone ends to hold the hydrogels in place within the defects. Hydrogels were crosslinked with 62.7 µmol or 67.5 µmol of a matrix metalloproteinase (MMP)-cleavable peptide sequence (GCRDVPMSMRGGDRCG) (Genscript Biotech Corporation, Piscataway, NJ, United States) to enable protease-mediated hydrogel degradation *in vivo*, washed 3 times with PBS, and loaded with 5 µg of BMP-2.

Unilateral femoral bone defects were created in 13-14 week old female Sprague Dawley rats (Charles River Laboratories) as previously described.^14^ Rats were anesthetized, administered analgesic, and aseptically prepared for surgery. A skin incision was made over the femur on the lateral aspect of the hind left leg, separation was created between the muscle groups, and the femur was isolated. A fixation plate designed to stabilize the femur after the bone defect^77^ was created was positioned onto the femur and held in place by a C-clamp. The plate was then secured to the femur via 4 screws. An oscillating surgical saw was used to create a 6-mm-wide, full thickness mid-diaphyseal bone defect. This defect is critically sized and does not heal without BMP-2 treatment.^57,76,78^ Hydrogels were placed into the defects. The surrounding musculature and the skin incision were closed in a three-layered approach using resorbable sutures and wound clips. Topical metronidazole paste was applied surrounding the surgical site to deter the animal from chewing at the incision site. Rats were observed postoperatively for any complications.

### 2.13. Radiographs and micro-CT

Longitudinal x-ray radiographs were taken at 4, 8, and 12 weeks after surgery (Faxitron MX20, 40 kV, 7 s). *In vivo* micro-computed tomography (micro-CT) was performed 6 weeks after surgery, and *ex vivo* micro-CT was performed after animals were euthanized at 12 weeks post-surgery. *In vivo* scans were performed using 48 μm voxel size, 55kVp voltage, 145 μA current, and 650 ms integration time (VivaCT 80, Scanco Medical, Brüttisellen, Switzerland). *Ex vivo* scans were performed using 24 μm voxel size, 55 kVp voltage, 145 μA current, and 750 ms integration time. Mineral was quantified within the central 5.5 mm of the defects. Scans were contoured beginning at the intact bone ends and to include either all new bone formed between bone ends or only bone within the defect space inside the diameter of the bone ends. Circular contours for defect bone were made to be 4.25 mm in diameter to encompass the intact femur ends. Investigators were blinded for all analyses.

### 2.14. Histology

Representative femurs from each group were selected for histological processing based on the mean micro-CT bone volume measurement for each group at 12 weeks. Femurs were simultaneously fixed and decalcified in Cal-Ex™ (Fisher Scientific) for 4 days at room temperature with shaking. The solution was replaced every two days and radiographs were taken every day to monitor decalcification. Femurs were then dehydrated in increasing concentrations of ethanol, cleared with xylene (Fisher Scientific), and infiltrated with paraffin wax. Samples were embedded in paraffin wax blocks and sectioned into 10 μm thick sections. Slides from each femur were stained using Hematoxylin & Eosin-y (H&E) (Abcam) for gross tissue morphology, Safranin-O/Fast Green (Saf-O) (Fisher Scientific) for proteoglycans in cartilage, and Picrosirius Red (Abcam) for collagen organization. Slides stained with H&E and Saf-O were imaged under the 5x brightfield scanning mode, and slides stained with Picrosirius Red were imaged using polarized light (Leica Thunder). Multiple images for each sample were stitched together using the Leica Las X automated multiple tile capture feature (LAS X).

### 2.15. Statistical Analysis

Compressive testing data were analyzed using MATLAB scripts to determine stiffnesses and yield points. SEM images were analyzed using ImageJ to determine fiber diameters and pore sizes. All data are plotted as mean and standard deviation. Statistically significant differences for each experiment were determined by performing a one-way ANOVA or two-way ANOVA as applicable with Tukey’s post-hoc test. A two-way ANOVA was performed on the micro-CT data to determine factor level effects. All analyses were performed using GraphPad Prism 10 (GraphPad Software).

## 3. Results

### 3.1. PEG-mal hydrogels can be synthesized within MEW scaffolds

Diamond-pore tubular scaffolds with a diameter of 5 mm and length of 5, 7.5, or 10 mm were printed using MEW on a custom AeroTech 3200 platform (**Figure 1A**).^52,79^ SEM images depict scaffolds with a diamond pore design (**Figure 1B**), mean fiber diameter of 29.45 ± 2.15 µm (**Figure 1C**), and mean pore area of 0.45 ± 0.03 mm^2^ (**Figure 1D**).

**Figure 1:**
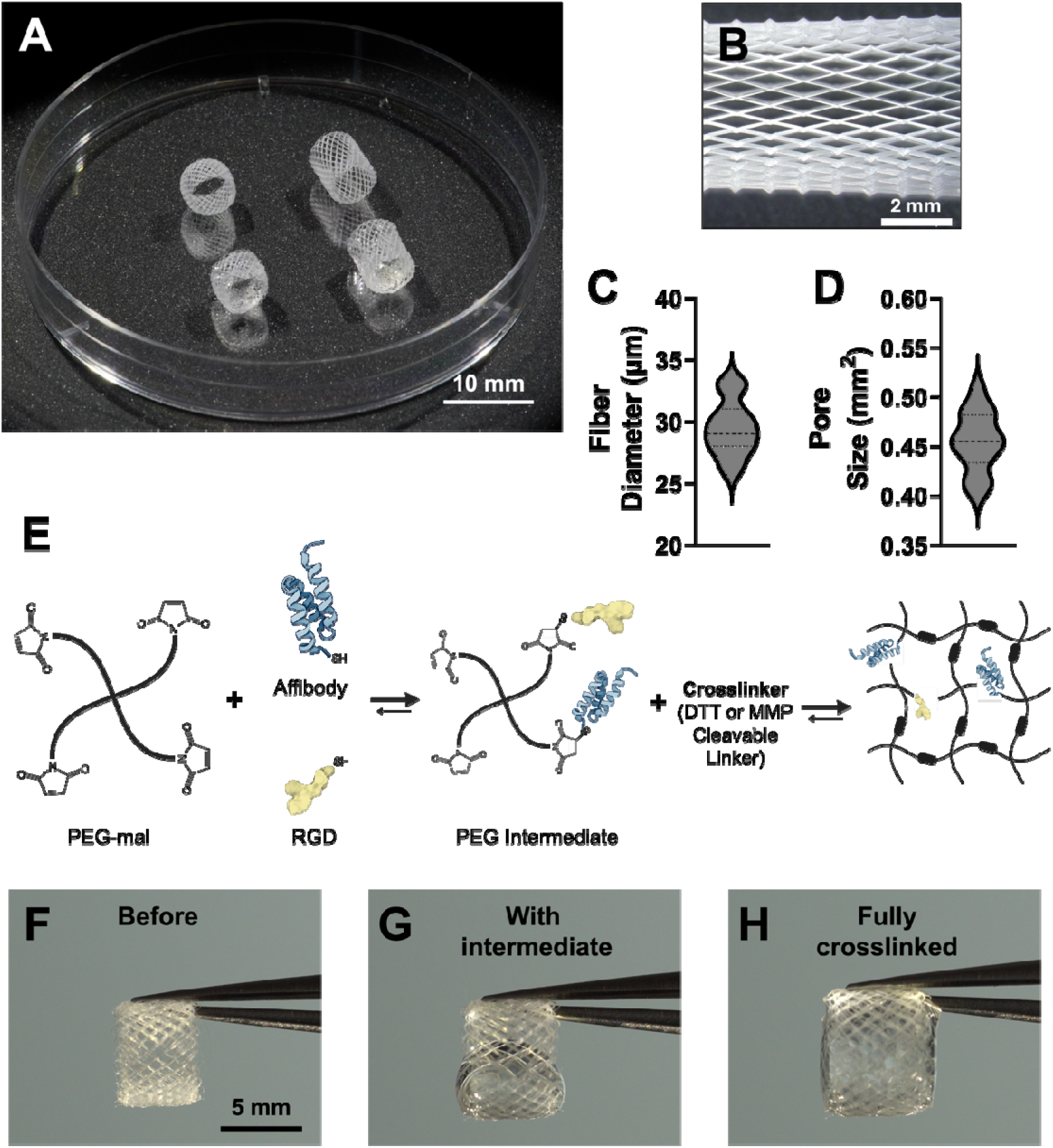
Fabrication of MEW scaffolds and scaffold-reinforced hydrogels. A) Photograph of tubular MEW scaffolds with lengths of 5 mm or 7.5 mm. Scale bar = 10 mm. B) SEM image of diamond pore design of MEW scaffold. Scale bar = 2000 μm. C) MEW scaffold fiber diameter distribution. D) MEW scaffold pore size distribution. E) Synthesis schematic for affibody- and RGD-conjugated PEG-mal hydrogels. A solution of PEG-mal, affibodies, and RGD peptides were mixed with either DTT or an MMP-cleavable peptide crosslinker to crosslink PEG-mal into hydrogels. F-H) Photos of fabrication of MEW scaffold-reinforced hydrogels. Scale bar = 5 mm. F) MEW scaffolds were suspended horizontally with tweezers. G) A solution of PEG-mal, affibodies, and RGD peptides and a solution of DTT or MMP-cleavable crosslinker were both pipetted into the tubular scaffold and rotated to allow the solution to mix. H) After mixing, the MEW scaffold was held horizontally for an additional 10 minutes to enable crosslinking of the PEG-mal hydrogel inside the tubular scaffold.

To synthesize hydrogels within MEW scaffolds, the PEG precursor solution (containing affibodies and/or RGD peptides) was slowly pipetted into a tubular scaffold held horizontally with tweezers (**Figure 1F**). The crosslinking solution containing either DTT or MMP-cleavable peptide crosslinker was added (**Figure 1G**), and the MEW scaffold was slowly rotated so that the solutions mixed and filled the inner space of the tube (**Figure 1H, Supplemental Video S1**). Within 10 seconds, the hydrogels crosslinked to a sufficient degree to remain within the scaffold. The scaffolds were suspended for an additional 10 minutes to ensure the DTT or peptide crosslinker could fully crosslink the available maleimides. The 5 mm long scaffolds were used to synthesize 100 µL hydrogels, and the 7.5 mm long scaffolds were used to synthesize 150 µL hydrogels. The conjugation of affibodies and RGD peptides to the maleimides on PEG-mal was not found to affect the mesh size of the hydrogels (**Figure S1**).

The effect of MEW scaffold reinforcement on hydrogel swelling and stability over 14 days *in vitro* was also assessed. Hydrogels with and without MEW scaffolds were initially swollen overnight (Day 0-1) and weighed and imaged periodically over 14 days. Hydrogels without MEW scaffolds visibly increased in size and changed shape after initial swelling, while reinforced hydrogels remained contained inside MEW scaffolds (**Figure S2A**). The weights of both the hydrogels with and without MEW scaffold reinforcement remained relatively consistent over 14 days after the initial swelling, at which point the hydrogels would typically be implanted *in vivo* (**Figure S2B**); however, the hydrogels without MEW scaffold reinforcement swelled more than the reinforced hydrogels.

### 3.2. MEW scaffolds improve hydrogel integrity without affecting stiffness

MEW scaffolds and hydrogels with or without MEW scaffolds and conjugated affibodies were subjected to radial compression testing to determine the effect of scaffold reinforcement on the mechanical properties of the constructs (**Figure 2A**). The MEW scaffold alone exhibited a stiffness of 96.4 ± 23.9 Pa, while the non-reinforced hydrogels displayed a stiffness of 159.1 ± 29.02 Pa without affibodies and 153.4 ± 34.8 Pa with affibodies (**Figure 2B**). The MEW scaffold-reinforced hydrogels displayed a stiffness of 210.1 ± 109.0 Pa without affibodies and 209.2 ± 55.73 Pa with affibodies. The results of compression testing suggest that MEW scaffold reinforcement did not impact the overall stiffness of the construct. At low compressive strain, the effects of the MEW scaffold were minimal; however, at 80% compressive strain, the reinforced hydrogels experienced a higher stress than the non-reinforced hydrogels and MEW scaffolds (**Figure 2C**), suggesting that MEW scaffold reinforcement begins to affect the amount of stress required to compress the hydrogels once they have been compressed by 80% of their original width. We also demonstrated that reinforced hydrogels were easier to manipulate with surgical forceps without damaging the hydrogel than non-reinforced hydrogels (**Supplemental Video S2**), which may make them easier to use in surgery than non-reinforced hydrogels.

**Figure 2:**
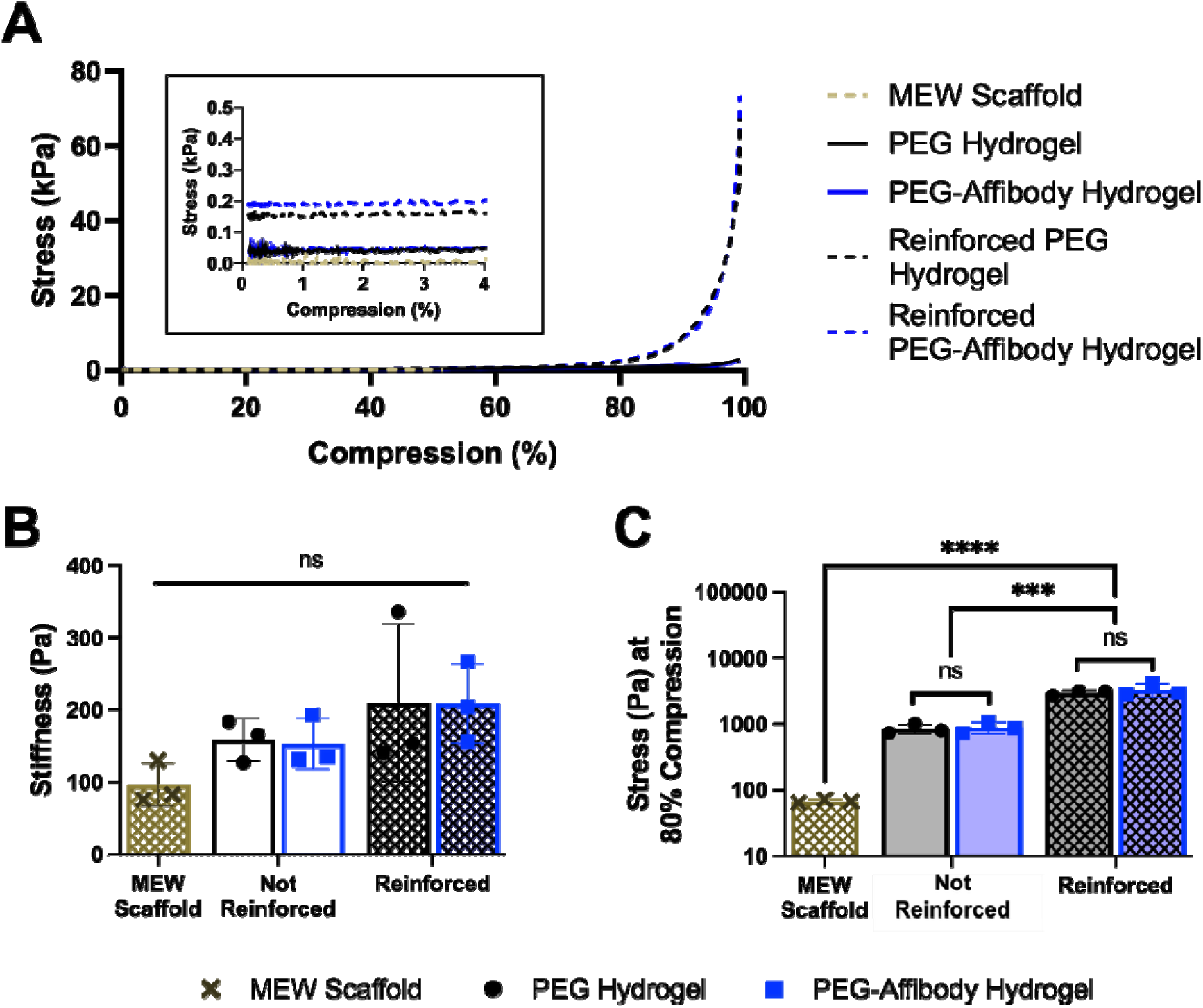
Compression testing of hydrogels with and without MEW scaffold reinforcement and conjugated affibodies. A) Stress vs. compressive strain of MEW scaffolds and hydrogels with or without MEW scaffold reinforcement and affibodies. Samples were compressed by up to 98% of their original width. Inlay plot demonstrates stress vs. compression at low compressive strain (0-4%), which was used for calculating stiffness. B) Stiffness of the MEW scaffold, hydrogels with and without affibodies, and scaffold-reinforced hydrogels with and without affibodies. C) Stress experienced by the MEW scaffold, hydrogels with and without affibodies, and scaffold-reinforced hydrogels with and without affibodies at 80% compressive strain. Statistical significance between groups was determined by one-way ANOVA with Tukey’s post-hoc test. n=3. *** p<0.001, **** p<0.0001, ns = not significant.

**Figure 3:**
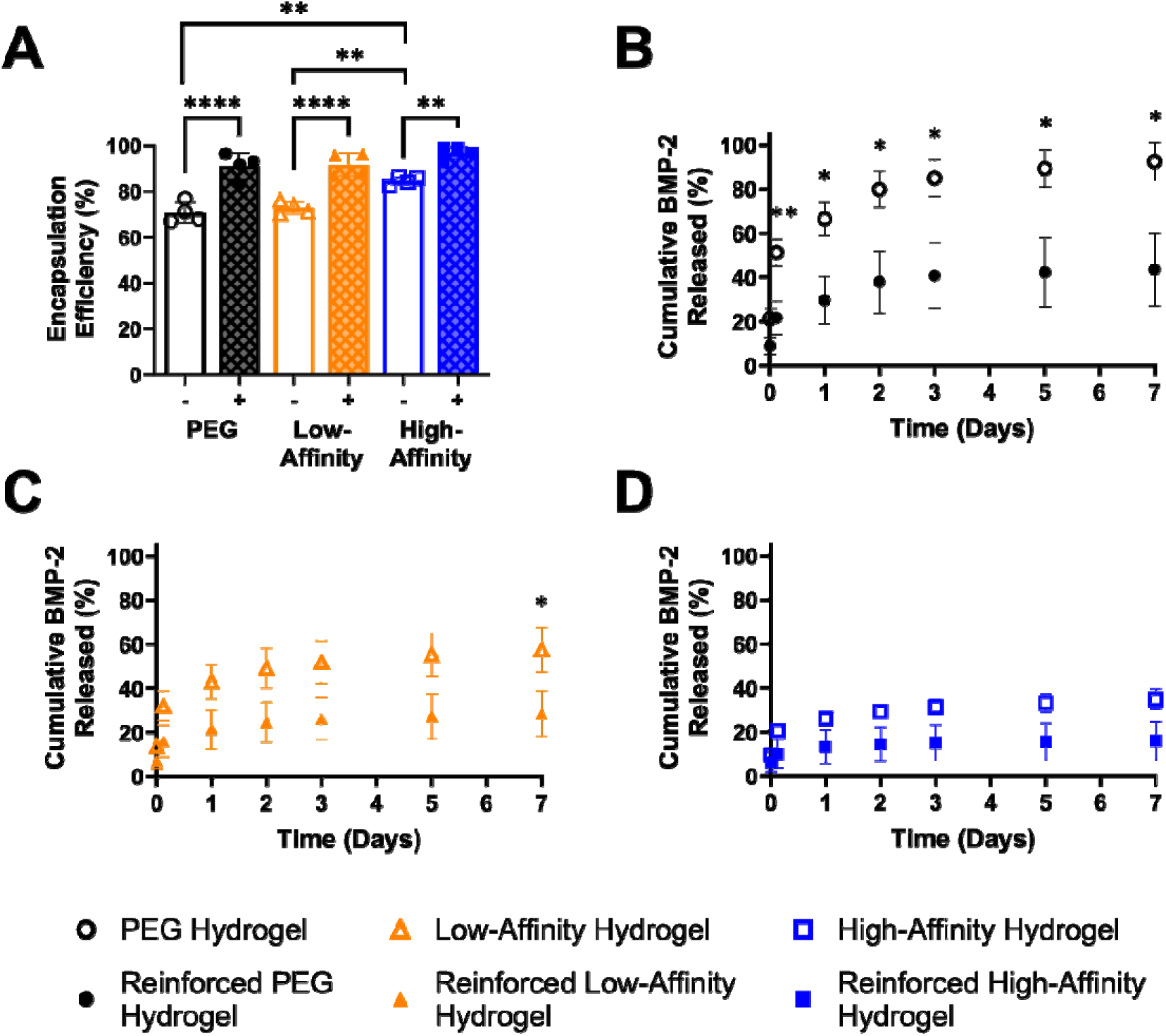
Effect of scaffold reinforcement on BMP-2 encapsulation and release. 100 ng of BMP-2 were loaded into PEG-mal hydrogels with or without MEW scaffold reinforcement and with or without conjugated low- or high-affinity BMP-2 affibodies. BMP-2 release into 10% (v/v) FBS in PBS was monitored over 7 days. A) Initial BMP-2 encapsulation efficiency as a function of affibody conjugation and scaffold reinforcement, where “–” indicates non-reinforced hydrogels and “+” indicates reinforced hydrogels. B-D) Release of BMP-2 from hydrogels over 7 days (B) without affibodies, (C) with low-affinity BMP-2-specific affibodies, and with (D) high-affinity BMP-2-specific affibodies. Statistical significance between groups at each time point was determined using two-way ANOVA with Tukey’ post-hoc test. n=3-4. * p<0.05, ** p<0.01, *** p<0.001, **** p<0.0001.

**Figure 4:**
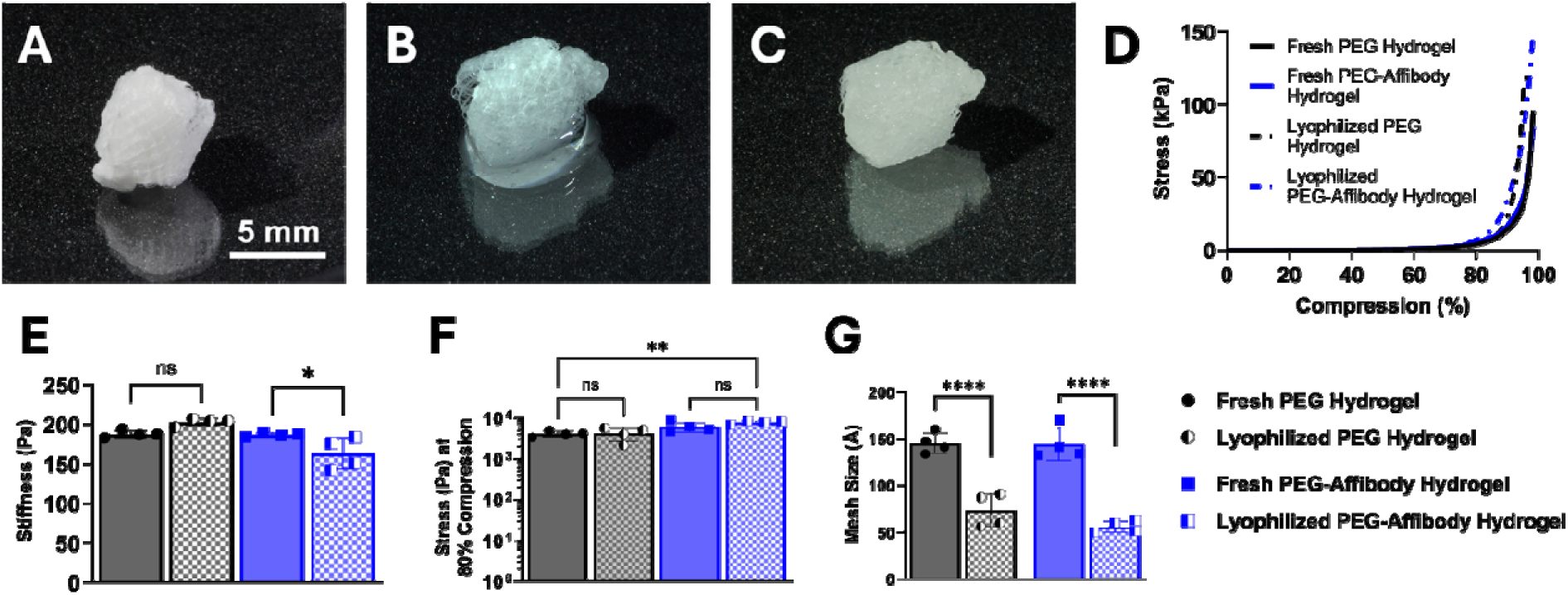
Effect of lyophilization and rehydration on physical properties of affibody-conjugated hydrogels. Fresh scaffold-reinforced hydrogels and scaffold-reinforced hydrogels after lyophilization and rehydration, with and without conjugated high-affinity affibodies, were evaluated. A) Lyophilized scaffold-reinforced hydrogel. Scale bar = 5 mm. B) Rehydration of lyophilized scaffold-reinforced hydrogel with approximately 130 µL of PBS at (B) 20 seconds and (C) 60 seconds. D) Stress vs. compressive strain of reinforced hydrogels. Samples were compressed up to 98% of their original width. Stress at low compressive strain (0-4%) was used to calculate stiffness. E) Stiffness of reinforced hydrogels. F) Stress experienced by reinforced hydrogels at 80% compressive strain. G) Mesh size of reinforced hydrogels calculated using Equilibrium Swelling Theory. Statistical significance between groups was determined using two-way ANOVA with Tukey’s post-hoc test. n=4. * p<0.05, ** p<0.01, **** p<0.0001, ns = not significant.

**Figure 5:**
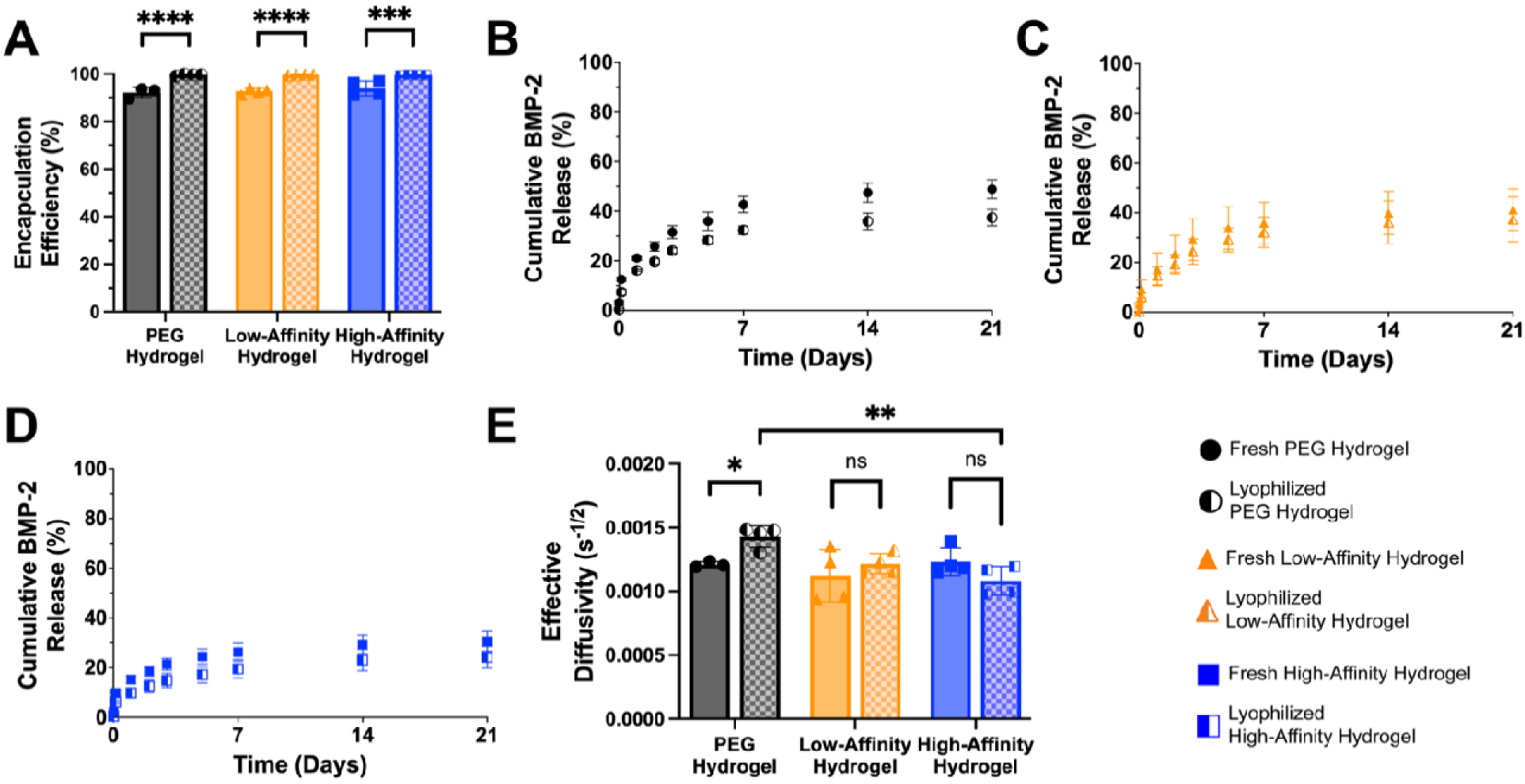
Effect of lyophilization and rehydration of scaffold-reinforced hydrogels on BMP-2 encapsulation and release. Fresh scaffold-reinforced hydrogels and scaffold-reinforced hydrogels after lyophilization and rehydration, with and without conjugated affibodies, were evaluated. 100 ng of BMP-2 were loaded into hydrogels, and BMP-2 release into 10% (v/v) FBS in PBS was monitored over 21 days. A) A) Initial BMP-2 encapsulation efficiency. B) BMP-2 release from hydrogels without affibodies. C) BMP-2 release from hydrogels containing low-affinity BMP-2-specific affibodies. D) BMP-2 release from hydrogels containing high-affinity BMP-2-specific affibodies. E) Effective diffusivity of BMP-2 from hydrogels calculated using the Korsmeyer-Peppas model. Statistical significance between groups at each time point was determined using two-way ANOVA with Tukey’s post-hoc test. n=3-4. * p<0.05, ** p<0.01, **** p<0.0001, ns = not significant.

**Figure 6:**
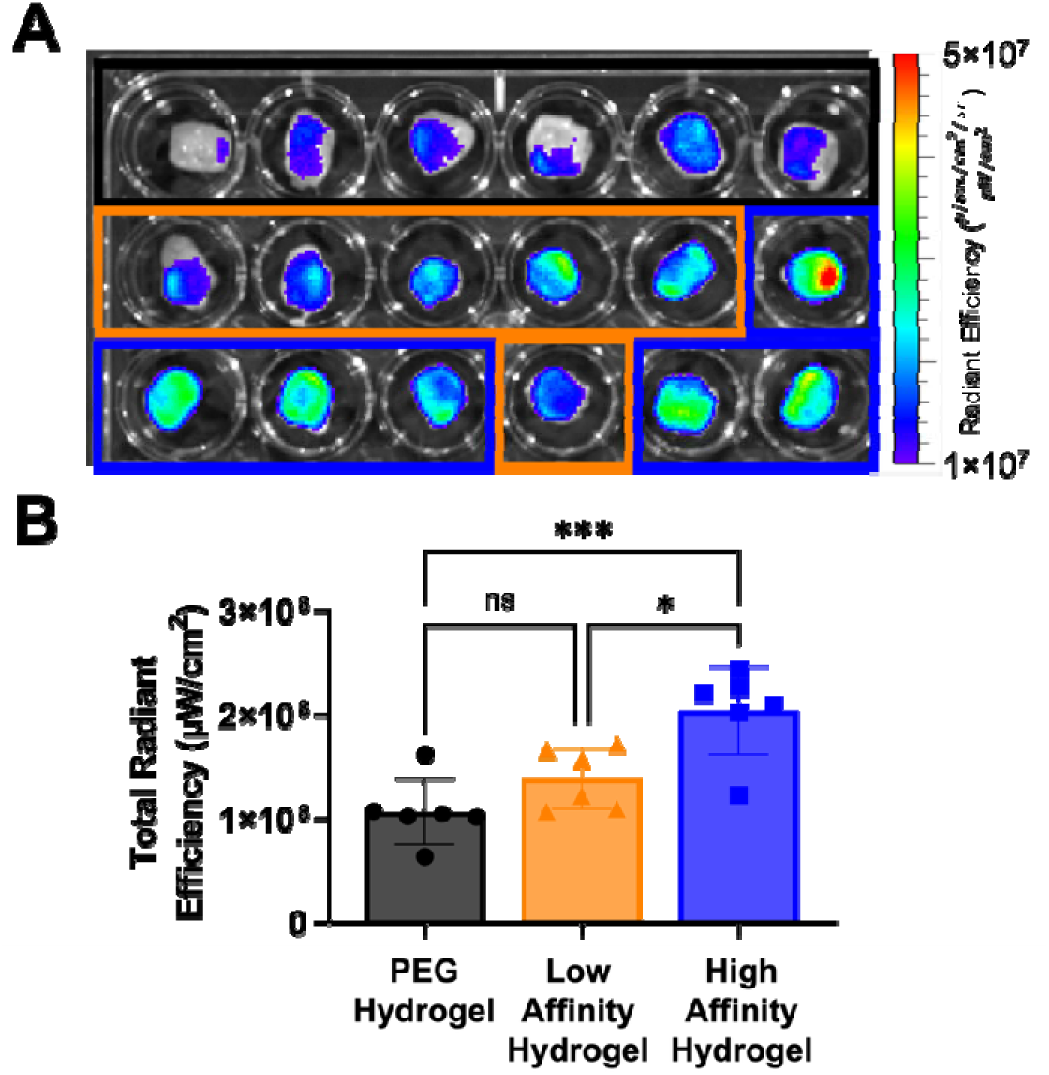
Retention of fluorescently labeled BMP-2 in scaffold-reinforced hydrogels after 3 weeks of subcutaneous implantation in rats. A) Photograph of excised hydrogels after 3 weeks with fluorescent signal overlay (excitation: 745 nm, emission: 800 nm). The top row, outlined in black, depicts hydrogel without affibodies, while hydrogels containing low-affinity BMP-2 affibodies and hydrogels containing high-affinity BMP-2 affibodies are outlined in orange and blue, respectively. B) Total radiant efficiency of the excised hydrogels. Statistical significance between groups was determined using one-way ANOVA with Tukey’s post-hoc test. n=6. * p<0.05, *** p<0.001, ns = not significant.

### 3.3. Effects of MEW scaffold reinforcement and affibodies on BMP-2 release

To investigate whether MEW scaffold reinforcement affects protein release from PEG-mal hydrogels, non-reinforced and reinforced hydrogels were loaded with BMP-2, and the initial encapsulation and subsequent release of BMP-2 from the constructs over 7 days were quantified by ELISA. Non-reinforced and reinforced hydrogels were fabricated without affibodies, with low-affinity BMP-2 affibodies, or with high-affinity BMP-2 affibodies. 100 μL non-reinforced hydrogels were formed as cylinders directly in the bottom of 2 mL centrifuge tubes with an approximate diameter of 10 mm and length of 4 mm such that a flat plane was formed. 100 μL reinforced hydrogels were fabricated within MEW scaffolds (5 mm diameter, 5 mm length) and placed in 2 mL centrifuge tubes for BMP-2 encapsulation and release. These different geometries and surface areas were unavoidable due to the non-reinforced PEG-mal hydrogels being unable to maintain their shape over time. As shown in **Figure S2A**, non-reinforced hydrogels quickly deformed during swelling. Similarly, when placed in a 2 mL microcentrifuge tube, these hydrogels would eventually take the shape of the tube. Thus, we chose to make these hydrogels at the bottom of the tube from the start of the experiment to avoid changes in geometry over time during protein release.

As we have previously demonstrated, high-affinity BMP-2 affibodies increased initial BMP-2 encapsulation in PEG-mal hydrogels compared to no affibodies and low-affinity BMP-2 affibodies (**Figure A**). MEW scaffold reinforcement also increased initial BMP-2 encapsulation compared to non-reinforced hydrogels, which could be attributed to both the increased exposed surface area of the reinforced hydrogel^39^ and potential adsorption of BMP-2 onto the PCL fibers.^80^ Scaffold reinforcement decreased BMP-2 release from hydrogels without affibodies at each time point over 7 days (**Figure B**). Conversely, scaffold reinforcement did not significantly impact BMP-2 release from hydrogels with affibodies except for a reduction in protein release with scaffold reinforcement of low-affinity affibody hydrogels at day 7 (**Figure C, 3D**). Similar to our previously reported results,^39^ significant differences in BMP-2 release were observed between all non-reinforced hydrogels at all time points after the initial release measurement, with hydrogels without affibodies releasing the most BMP-2, low-affinity hydrogels releasing less BMP-2 than the hydrogels without affibodies, and high-affinity hydrogels releasing the least BMP-2 (**Figure S3A**). Reinforced hydrogels without affibodies released significantly more BMP-2 than the hydrogels with high-affinity affibodies at all time points after day 1 (**Figure S3B**). These data demonstrate that both protein-material affinity interactions and scaffold reinforcement decrease BMP-2 release. Additionally, BMP-2 release from reinforced PEG-mal hydrogels with and without high-affinity BMP-2 affibodies was compared to release from collagen sponges as the clinical standard (**Figure S4**). Collagen sponges encapsulated the highest amount of BMP-2, although all constructs encapsulated >88% of the loaded BMP-2 (**Figure S4A**). There were no significant differences between the amount of BMP-2 released from reinforced PEG-mal hydrogels and collagen sponges at any time point (**Figure S4B**). In contrast, collagen sponges released a significantly higher amount of BMP-2 than reinforced hydrogels with affibodies at each time point after 6 hours, again underscoring the importance of affinity interactions in controlling BMP-2 release.

Cumulative BMP-2 release data were fitted to the Korsmeyer-Peppas drug release model (**Equation 3**) to determine the mechanism of BMP-2 release from the hydrogels. The diffusion exponent, n, which defines the mechanism of drug release, was 0.24 ± 0.03 for non-reinforced hydrogels and 0.28 ± 0.05 for reinforced hydrogels (**Figure S5A, S5B**). These values fall below the upper bound for Fickian diffusion (n ≤ 0.50 for slab geometries for the non-reinforced hydrogels, and n ≤ 0.45 for cylindrical geometries for the reinforced hydrogels),^81^ indicating Fickian diffusion as the mechanism of protein release. By fixing the n exponent to its upper bound (0.5 for non-reinforced hydrogels, and 0.45 for reinforced hydrogels), the effective diffusivity K could be determined from **Figures S5C and S5D** and compared between geometries. In both non-reinforced and reinforced hydrogels, the high-affinity affibody hydrogels displayed a reduced effective BMP-2 diffusivity (i.e., release rate) compared to the low-affinity affibody and PEG-mal hydrogels (**Figure S5E, S5F**), aligned with previously observed responses.^39,59^

### 3.4. Scaffold reinforcement improves hydrogel integrity during lyophilization and rehydration

To determine the effect of lyophilization on hydrogel properties, hydrogels with and without MEW scaffold reinforcement and conjugated affibodies were immediately frozen and lyophilized following fabrication. While hydrogels without MEW scaffolds collapsed and did not retain their shape during the lyophilization process, the reinforced hydrogels retained their shape within the MEW scaffold and could be easily handled (Error! Reference source not found.**A**). Lyophilized hydrogels were rehydrated with PBS (Error! Reference source not found.**B**) and found to uptake approximately 130 µL of liquid within one minute (**Supplemental Video S3**), while remaining confined within the MEW scaffold (Error! Reference source not found.**C**).

Fresh scaffold-reinforced hydrogels and hydrogels that had undergone lyophilization and rehydration were compressed by 98% strain to determine construct stiffness (**Figure D**). The stiffness of the reinforced hydrogels without affibodies was not affected by lyophilization and rehydration, but the stiffness of the reinforced affibody-conjugated hydrogels was reduced from 186 ± 3 Pa before lyophilization to 163 ± 20 Pa after lyophilization and rehydration (Error! Reference source not found.**E**). Lyophilization and rehydration did not affect the stress experienced by the hydrogels at 80% compression (Error! Reference source not found.**F**). Equilibrium Swelling Theory was used to determine the mesh size of the reinforced hydrogels with and without affibodies and before and after lyophilization and rehydration. Rehydrated lyophilized hydrogels exhibited a significant reduction in mesh size compared to fresh hydrogels, going from approximately 140 Å to approximately 60 Å (**Figure G**). These results indicate that the crosslinked polymer network of the hydrogels was disrupted by the lyophilization and rehydration process, which has also been demonstrated in other studies.^19,55,82^ Lyophilization may have caused an irreversible collapse of the crosslinked polymer network, such that rehydration could not restore its original porosity. Regardless, the MEW scaffold reinforcement maintained the integrity and bulk mechanical properties of the hydrogel to ensure it could still be adequately handled during processing. The MEW fibers themselves did not appear to be affected by freezing and lyophilization due to the low water content of PCL, low second moment of area, and thermal stability of the polymer at low temperatures used for freezing before lyophilization.^83,84^

### 3.5. Effects of lyophilization and rehydration of reinforced hydrogels on BMP-2 release

To determine the effect of lyophilization and rehydration of scaffold-reinforced hydrogels on BMP-2 release, we loaded fresh and lyophilized hydrogels with 100 ng BMP-2 for 30 minutes, which is similar to the amount of time that BMP-2 solution is allowed to rehydrate the absorbable collagen sponges used in the clinic prior to their use in surgery.^71^ Fresh, hydrated hydrogels were loaded with a small volume (20 µL) of 5 µg/mL BMP-2 in PBS because they were already water-swollen, while lyophilized hydrogels were rehydrated with 130 µL of 0.769 µg/mL BMP-2 in PBS. Lyophilized and rehydrated hydrogels encapsulated more BMP-2 regardless of affibody conjugation (**Figure A**). No differences in cumulative BMP-2 release were observed between fresh hydrogels and lyophilized and rehydrated hydrogels without affibodies (**Figure B**), with low-affinity BMP-2 affibodies (**Figure C**), or with high-affinity BMP-2 affibodies (**Figure D**). Regardless of lyophilization, all hydrogels containing high-affinity BMP-2 affibodies released significantly less BMP-2 than hydrogels without affibodies (p = 0.0051 on day 7 for fresh hydrogels and p = 0.0138 on day 7 for lyophilized hydrogels, measured by 2-way ANOVA with Tukey’s post-hoc test, n=3-4), which is consistent with previous results.^39,59^ These data suggest that affibody-conjugated hydrogels retain their ability to control BMP-2 release following lyophilization and rehydration. BMP-2 release profiles were fit to the Korsmeyer-Peppas model (**Equation 3**) to determine the effective diffusivity of BMP-2 through each hydrogel formulation, revealing that only lyophilization and rehydration of the hydrogels without affibodies increased BMP-2 diffusivity (**Figure E**). Similar to previous results,^39,59^ effective BMP-2 diffusivity was lower in hydrogels containing high-affinity BMP-2 affibodies compared to hydrogels without affibodies. This result further supports the theory that affibodies remain functional and retain their ability to control BMP-2 release even after lyophilization and rehydration of the construct and that BMP-2-affibody affinity influences BMP-2 release rather than hydrogel mesh size.

### 3.6. High-affinity BMP-2 affibodies increase BMP-2 retention within hydrogels implanted *in vivo*

Fluorescently labeled BMP-2 was loaded into 150 µL scaffold-reinforced hydrogels with or without conjugated affibodies and implanted in the dorsal subcutaneous space of Sprague Dawley rats to investigate the effect of affibodies on local BMP-2 retention *in vivo*. No adverse effects were observed post-operatively. After 3 weeks, the rats were euthanized, and the hydrogels were explanted, photographed, and fluorescently imaged (**Figure A**). Quantification of fluorescent signal revealed that hydrogels containing high-affinity BMP-2 affibodies retained significantly more BMP-2 than both hydrogels without affibodies and hydrogel containing low-affinity BMP-2 affibodies (**Figure B**). X-ray radiographs revealed no mineral formation in the hydrogels after 3 weeks.

3.7. Effects of scaffold reinforcement and high-affinity affibodies on BMP-2-induced bone formation in rat femoral bone defects

Since high-affinity BMP-2 affibodies significantly increased BMP-2 retention in hydrogels implanted in the subcutaneous space, we next sought to investigate the effects of affibody conjugation and scaffold reinforcement on the efficacy of PEG-mal hydrogels as BMP-2 delivery vehicles for bone regeneration. We implanted PEG-mal hydrogels without affibodies or scaffold reinforcement, scaffold-reinforced hydrogels without affibodies, and scaffold-reinforced hydrogels with high-affinity BMP-2 affibodies in 6-mm femoral defects in Sprague Dawley rats. All hydrogels were loaded with 5 µg of BMP-2, which has been shown to stimulate bone bridging across the defect space within 12 weeks in this model.^57,85^ Representative longitudinal radiographs at 4, 8, and 12 weeks demonstrate progressive mineralization throughout the defect space in all treatment groups (**Figure 7**). Based on radiographs at 12 weeks post-surgery, bone bridging was observed in 9 out of 11 rats treated with PEG-mal hydrogels without affibodies or scaffold reinforcement, 12 of 12 rats treated with scaffold-reinforced hydrogels without affibodies, and 12 of 12 rats treated with scaffold-reinforced hydrogels with high-affinity BMP-2 affibodies.

**Figure 7:**
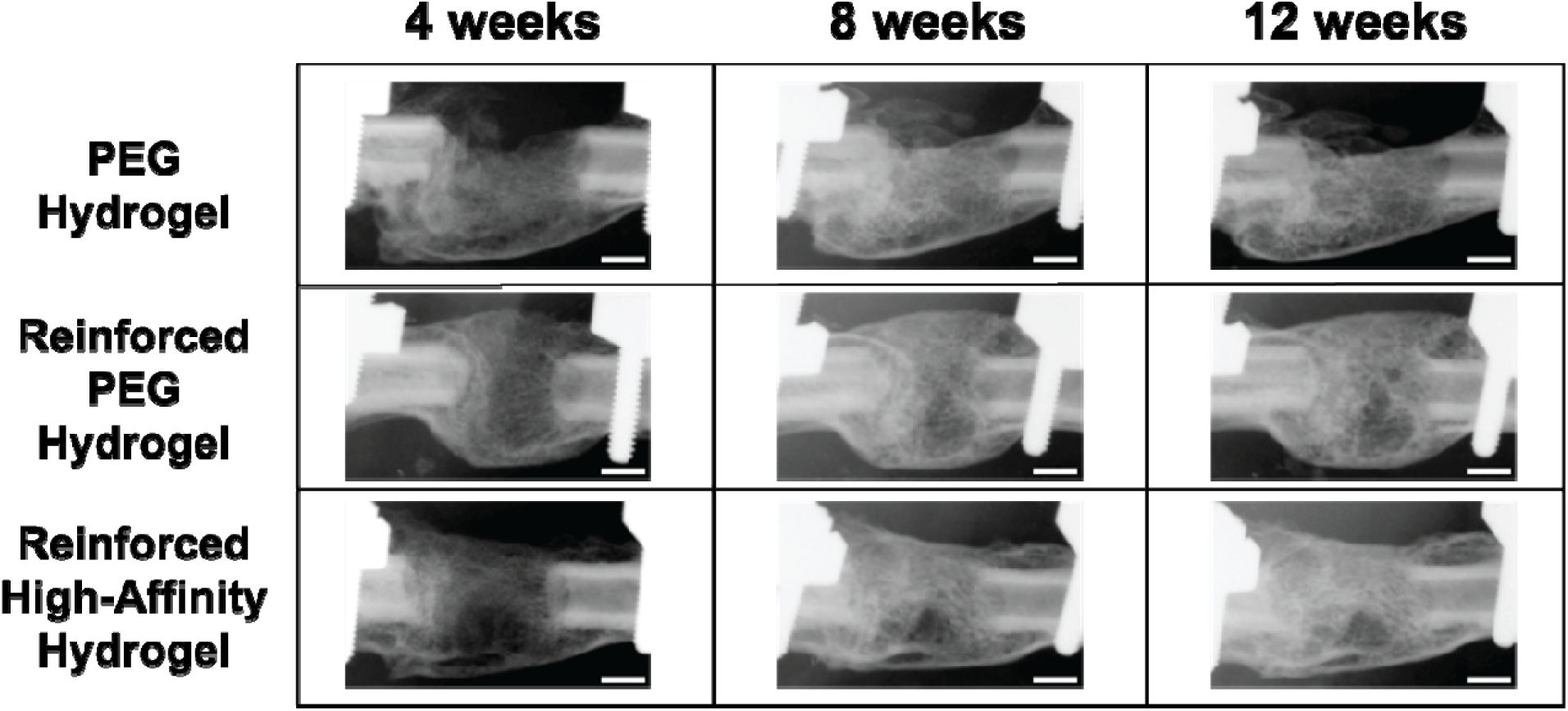
Representative longitudinal radiographs of rats treated with BMP-2-loaded hydrogel with or without scaffold reinforcement and high-affinity BMP-2 affibodies. 6-mm full-thickness femoral defects were treated with 5 µg of BMP-2 delivered in PEG-mal hydrogels, hydrogels reinforced with MEW scaffolds, and hydrogels reinforced with MEW scaffolds and containing high-affinity BMP-2 affibodies. Longitudinal *in vivo* radiographs were taken at 4, 8, and 12 weeks post-surgery. Scale bars = 2 mm.

Micro-CT was performed *in vivo* at 6 weeks and *ex vivo* at the conclusion of the study at 12 weeks to quantify total bone volume and bone volume inside the defect space. Two-way ANOVA factor analysis revealed that MEW scaffold reinforcement significantly increased total and defect bone volume relative to non-reinforced hydrogels at 6 weeks (**Figure 7A**) and total bone volume relative to non-reinforced hydrogels at 12 weeks (**Figure 7E**). The presence of high-affinity BMP-2 affibodies did not affect bone volume at either time point. Representative mid-sagittal sliced micro-CT reconstructions for the hydrogels without affibodies or reinforcement (**Figure 8B**), scaffold-reinforced hydrogel without affibodies (**Figure 8C**), and scaffold-reinforced affibody-conjugated hydrogels (**Figure 8D**) at 6 weeks were consistent with radiographs depicting new bone formation throughout the defect space for all treatment groups. Representative mid-sagittal sliced micro-CT reconstructions of the same animals at 12 weeks demonstrated similar bone morphology for all treatment groups (**Figure 8F-H**).

**Figure 8:**
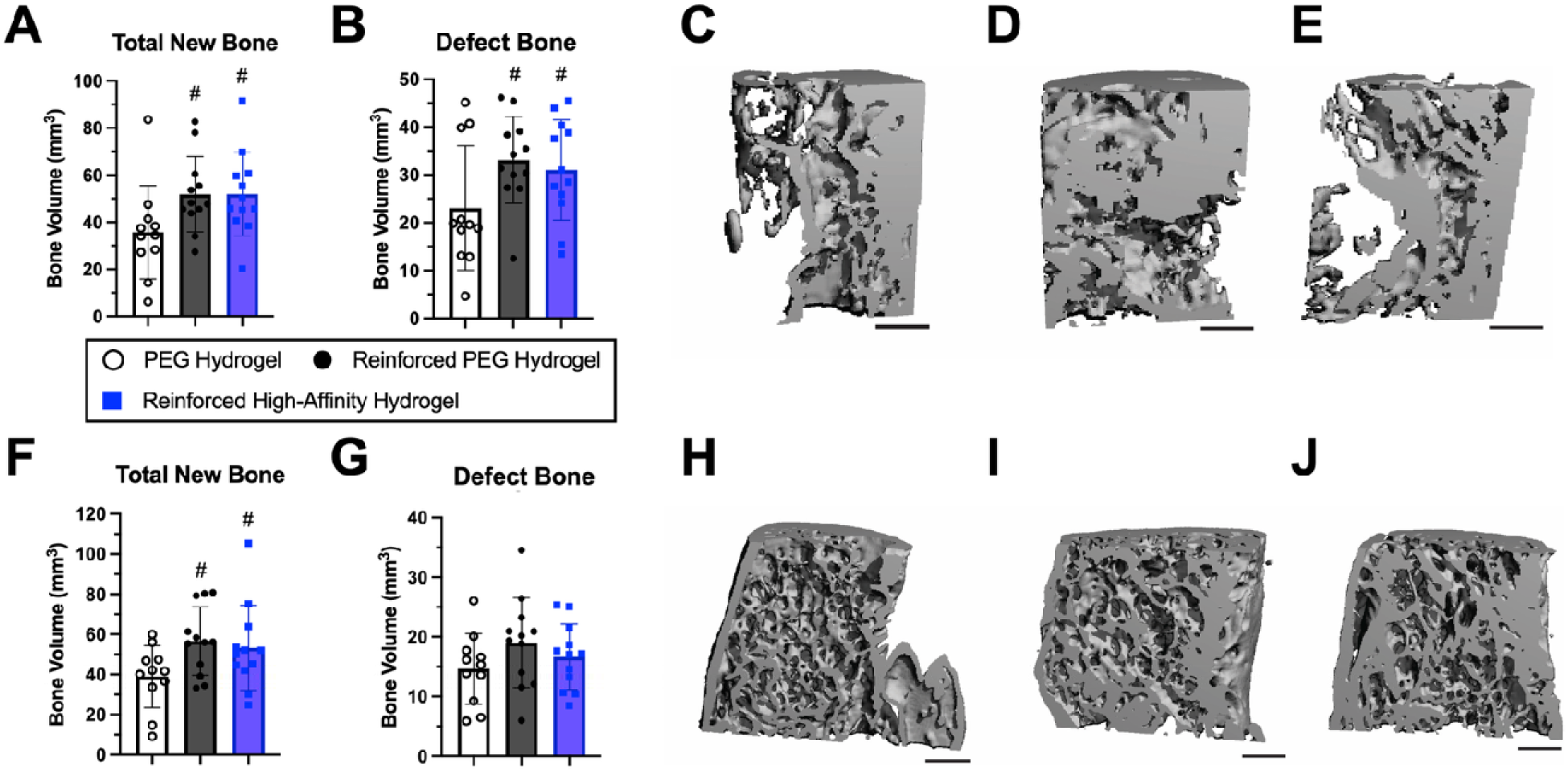
Micro-CT quantification and 3D reconstructions of regenerated bone at 6 and 12 weeks post-surgery. 6-mm full-thickness femoral defects were treated with 5 µg of BMP-2 delivered in PEG-mal hydrogels, hydrogels reinforced with MEW scaffolds, and hydrogels reinforced with MEW scaffolds and containing high-affinity BMP-2 affibodies. A) *In vivo* micro-CT quantification of total new bone and bone formed within the defect margins for each treatment group after 6 weeks. Representative mid-sagittal sliced micro-CT reconstructions for B) hydrogels without affibodies or reinforcement, C) scaffold-reinforced hydrogel without affibodies, and D) scaffold-reinforced affibody-conjugated hydrogels after 6 weeks. E) Ex vivo micro-CT quantification of total new bone and defect bone formed for each treatment group after 12 weeks. Representative mid-sagittal sliced micro-CT reconstructions for F) hydrogels without affibodies or reinforcement, G) scaffold-reinforced hydrogel without affibodies, and H) scaffold-reinforced affibody-conjugated hydrogels after 12 weeks. Scale bars = 1 mm. Statistical significance between groups was determined using two-way ANOVA with Tukey’s post-hoc test for factor effect analysis. n=11-12. # factor effect of MEW scaffold reinforcement.

### 3.8. Histological analysis

Histological staining was performed on representative femurs isolated after 12 weeks post-surgery. Hematoxylin and eosin-y (H&E) staining revealed similar cellularity and morphological features for all treatment groups (**Figure 9A-C**). In defects treated with reinforced hydrogels, the MEW fiber scaffolds can be observed in the H&E-stained sections as small gaps in the tissue (yellow arrows); however, the fibers are surrounded by tissue and do not appear to have impacted tissue infiltration and bone formation in the defect space. Safranin O staining revealed minimal proteoglycan staining in all defects (**Figure 9D-I**). Histological slides stained with Picrosirius Red and imaged under polarized light to visualize collagen revealed thicker, packed yellow-orange fibers, which are often associated with collagen type I, in defects treated with the non-reinforced PEG-mal hydrogels (**Figure 9J, 9M)**. The reinforced hydrogels with and without affibodies both displayed thinner fibers, which are typically associated with collagen type III (yellow-green, indicated with blue arrows), potentially indicating lamellar bone^86,87^ (**Figure 9K-L, 9N-O**).

**Figure 9:**
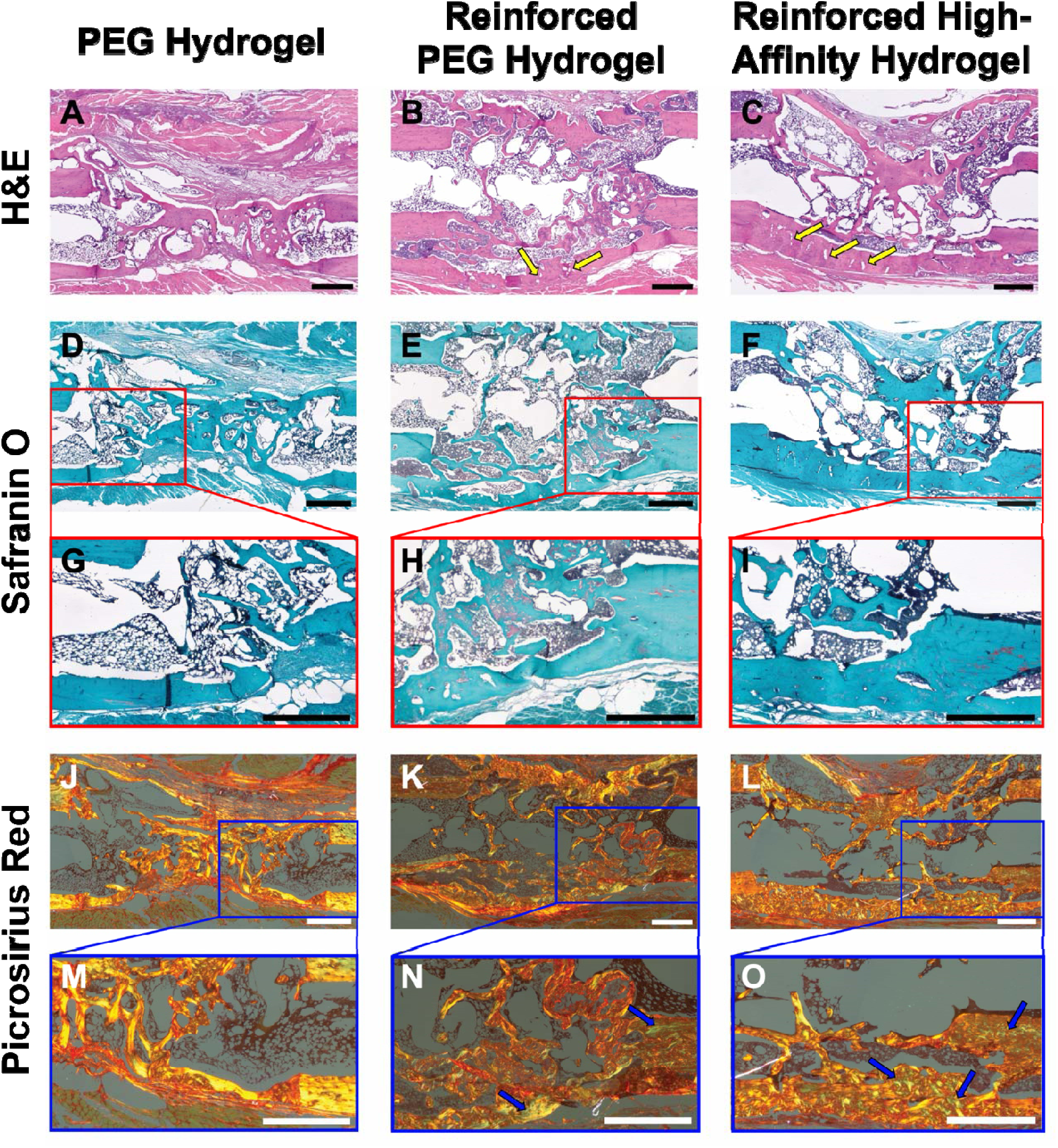
Histological staining of femoral bone defects at 12 weeks post-surgery. 6-mm full-thickness femoral defects were treated with 5 µg of BMP-2 delivered in PEG-mal hydrogels with or without MEW scaffold reinforcement and high-affinity BMP-2 affibodies. A-C) Hematoxylin and eosin-y staining of representative femurs. Yellow arrows indicate holes in tissue from MEW scaffold. D-F) Safranin O staining of representative femurs. G-I) Zoomed-in images of Safranin O staining. J-L) Picrosirius Red staining imaged under polarized light of representative femurs. M-O) Zoomed-in images of Picrosirius Red staining. Blue arrows indicate yellow-green staining. Scale bars = 1 mm.

## 4. Discussion

Recombinant BMP-2 delivery has demonstrated success in treating severe bone injuries that require surgical intervention.^3,11^ However, limitations in the application and overall efficacy of BMP-2 treatment via the clinical collagen sponge still exist due to its propensity to cause side effects associated with poor BMP-2 localization, resulting in soft tissue swelling and heterotopic ossification.^3^ Developing clinically relevant recombinant protein delivery vehicles for bone regeneration remains a challenge, as hydrogels must provide sustained protein delivery to mimic the well-coordinated bone healing cascade, support cell and tissue infiltration and subsequent mineralization, withstand surgical implantation and biologically relevant stresses, and have the potential to be shelf-stable to broaden accessibility. We developed a MEW scaffold-reinforced PEG-mal hydrogel that controlled the delivery of BMP-2 via protein-material affinity interactions, could be easily handled for surgical implantation, and was amenable to lyophilization and rehydration for extended shelf stability, increasing its potential for facile clinical adoption. We demonstrated that affibody-conjugated PEG-mal hydrogels could be easily fabricated within tubular MEW scaffolds without impacting their stiffnesses or affinity-controlled BMP-2 release. Scaffold reinforcement also enabled the lyophilization and rehydration of PEG-mal hydrogels without disintegration. Despite a significant change in mesh size, lyophilized and rehydrated affibody-conjugated hydrogels retained the ability to control BMP-2 release through protein-material affinity interactions. *In vivo*, affibody conjugation enhanced BMP-2 retention in subcutaneous implants, while MEW scaffold reinforcement increased total bone volume and defect bridging in rat femoral bone defects.

Given the delicate nature of most hydrogel- and sponge-based protein delivery vehicles, scaffold reinforcement presents a promising strategy to improve hydrogel integrity and handling for clinical use. We and others have demonstrated that combining hydrogels with MEW fiber scaffolds can leverage the biochemical and biophysical properties of both types of materials.^43,88,89^ Here, we leveraged the surface tension between the PEG-mal hydrogel precursors and PCL tubular scaffolds to generate highly reproducible affibody-conjugated PEG-mal hydrogels confined within the scaffold. This innovation overcame the challenge of inconsistent hydrogel sizes when hydrogels were synthesized in molds. MEW scaffold reinforcement has also been used by others to improve hydrogel mechanical properties and surgical use. Visser, et al. demonstrated that infusing gelatin methacrylate or alginate hydrogels into MEW fiber scaffolds with a 3D box structure increased the stiffness of the composite constructs by 54-fold and 15-fold, respectively.^43^ Similarly, Eicholz, et al. combined an outer tube fabricated using fused deposition modeling with a box structure of MEW microfibers to generate cylindrical scaffolds with precise inner micro-architecture to guide bone regeneration.^90^ These constructs provided sustained BMP-2 delivery and supported bone repair in a rat femoral defect; however, bone bridging was not observed in all animals, and the addition of BMP-2 increased heterotopic bone formation.

In both cases, these scaffold designs introduced MEW fibers throughout the entire construct, unlike our scaffold reinforcement strategy, in which a hollow tube of MEW fibers encircled a bulk hydrogel. While the goal of these other designs was to increase construct stiffness to provide mechanical stability to tissue defects, we designed a tissue engineered construct for use in a mechanically stabilized long bone defect, and thus the purpose of our design was to instead improve construct handling during surgery while maintaining the original hydrogel stiffness and without impacting affinity-controlled BMP-2 release or subsequent tissue infiltration *in vivo*. The lower stiffness of our constructs is likely to be more advantageous for bone repair in defects with robust mechanical fixation, which can be enhanced by using softer scaffolds.^17,18^ Furthermore, given the relatively slow *in vivo* degradation rate of microfiber PCL,^91,92^ the low PCL fiber content of our scaffold design may enable more robust bone remodeling over longer periods of time. It should be noted that our scaffold reinforcement strategy was only shown to enhance hydrogel integrity under radial compressive forces, such as those expected during surgical handling for implantation and exerted by adjacent soft tissue in the injury site. Since our bone defect model involved mechanical stabilization along the axis of the bone, our hydrogels were stress-shielded and not expected to experience significant axial compressive forces.^77,93^ Additional engineering may be required to design a robust material that resists axial compression or deformation under tension or shear forces. Although other scaffold geometries were not tested in this study, MEW scaffold mechanical properties can be widely tailored using a variety of the printing patterns and scaffold geometries.^43,52^ While the current MEW scaffold design is not intended to be load-bearing, further optimization could be performed to increase its stiffness for load-bearing applications.^43^

An additional advantage of reinforcing our hydrogels with MEW scaffolds was that we were able to flash freeze, lyophilize, and rehydrate the constructs while retaining their original structure and mechanical properties. We demonstrated that the constructs could be rapidly rehydrated with a highly concentrated protein solution, in a manner similar to the absorbable collagen sponge clinically used for BMP-2 delivery.^94^ Aside from a small decrease in the stiffness of the lyophilized and rehydrated affibody-conjugated hydrogel compared to the non-processed hydrogel, hydrogel mechanical properties remained largely unchanged before and after lyophilization and rehydration. Our results corroborate other work in the field that has demonstrated that lyophilization either has no effect or minimally decreases the mechanical properties of hydrogels.^55,95^ Interestingly, lyophilized and rehydrated hydrogels without affibodies released BMP-2 at a faster rate than non-processed hydrogels, despite a significant decrease in hydrogel mesh size, which is typically expected to hinder protein diffusion through the hydrogel network.^22,67^ However, few studies have been conducted to determine the effect of lyophilization on diffusion-controlled protein release,^96,97^ and none have been conducted to determine the effect of lyophilization on affinity-controlled protein release. Our rehydrated hydrogels containing high-affinity BMP-2 affibodies released less protein at a slower rate than hydrogels without affibodies, indicating that affibody-conjugated hydrogels retain the BMP-2-binding capacity necessary to control BMP-2 release. Thus, we show, for the first time, that protein-material affinity interactions mediated by affibodies can be retained post-lyophilization and rehydration. These promising results open the doors to further investigation of lyophilization and rehydration of affibody-conjugated hydrogels as a strategy to increase their shelf stability. Our future work will examine the function of rehydrated affibody-conjugated hydrogels for affinity-controlled release over longer periods of time post-lyophilization and under accelerated aging conditions.

While the affinity interactions between BMP-2 and the BMP-2-specific affibodies play a critical role in controlling BMP-2 release from these hydrogels, BMP-2 release is also dependent on other properties of the hydrogel, such as mesh size, swelling, and stiffness, which may be influenced by polymer and crosslinker choice, adhesive peptide conjugation, and crosslinking density. The highest BMP-2 loading used in these studies was 5 µg per hydrogel, which was used in the animal studies. This loading required the use of 96.2 nmol affibodies, which occupied 2.5% of the total available maleimides. While many more maleimides are available on 4-arm PEG-mal even at this highest BMP-2 dose, the maleimides are also required for conjugating cell adhesive peptides (RGD) and crosslinking with DTT or peptide crosslinker. As such, increasing the affibody content in a 5% (w/v) 4-arm PEG-mal hydrogel will reduce the available maleimides for crosslinking and adhesive peptide conjugation, thereby decreasing the stiffness and increasing the swelling and mesh size of the hydrogel.^98–100^ Thus, maximizing affibody concentration in the hydrogels would not necessarily maximize BMP-2 loading and minimize BMP-2 release. Substituting 4-arm PEG-mal with 8-arm PEG-mal could provide more maleimides for crosslinking and affibody/peptide conjugation and has been shown to increase hydrogel stiffness while decreasing swelling and gelation time.^100^ Thus, 8-arm PEG-mal could be used to increase the affibody content of the hydrogel while still allowing robust crosslinking and adhesive peptide conjugation to retain the hydrogel’s mechanical properties and permissiveness to cell infiltration.

Although several studies have demonstrated that other types of natural and engineered protein-material affinity interactions can increase protein retention in biomaterials implanted *in vivo*,^14,57,76,101^ this is the first study to directly demonstrate that protein-specific affibodies can increase protein retention *in vivo*. PEG-mal hydrogels containing high-affinity BMP-2 affibodies retained significantly more BMP-2 subcutaneously after 3 weeks *in vivo* compared to hydrogels without affibodies or with low-affinity BMP-2 affibodies, motivating further investigation in a pre-clinical model of bone repair. We chose to investigate bone regeneration in a well-characterized, critically-sized bone defect in the femur of a rat, which is amenable to bulk hydrogel implantation, longitudinal imaging, and mechanical testing and demonstrates a dose-dependent response to sustained BMP-2 delivery.^14,57,58^ 5 µg of BMP-2 were delivered in PEG-mal hydrogels, PEG-mal hydrogels reinforced with MEW scaffolds, and PEG-mal hydrogels reinforced with MEW scaffolds and containing high-affinity BMP-2 affibodies. This dose of BMP-2 has been shown to stimulate bone bridging in this model,^57,58^ and similar PEG-mal hydrogels crosslinked with MMP-cleavable crosslinkers have been shown to effectively deliver BMP-2 in other rodent models of long bone defects.^102^ MEW scaffold reinforcement increased total new bone volume at 12 weeks and both total new bone and orthotopic bone volume at 6 weeks, demonstrating that improving containment of the PEG-mal hydrogel in the bone defect could enhance bone repair. However, despite improving BMP-2 retention within the implantation site, the incorporation of high-affinity BMP-2 affibodies did not significantly impact bone volume or morphology over 12 weeks, corroborating other results that demonstrate that improved BMP-2 retention may not necessarily result in improved bone healing in this injury model.^76^ This result may be due to the fact that the 5 µg BMP-2 dose was high enough to robustly regenerate bone in this model without the need for affinity-controlled protein retention, while variable bone regeneration has been observed in this model with lower BMP-2 doses.^57^ Histological analyses revealed that the MEW scaffolds did not interfere substantially with tissue infiltration, and defects treated with reinforced hydrogels also displayed more organized lamellar bone compared to defects treated with non-reinforced PEG-mal hydrogels.

Solid electrospun PCL nanofiber mesh tubes have been similarly used to contain alginate hydrogels delivered in this femoral defect model.^14,57,58^ In a previous study, the addition of 1-mm perforations in the mesh tubes, accounting for 10% of the total tube surface area, was found to accelerate early bone growth, likely through the facilitation of early cell and tissue infiltration.^58^ Building upon this idea, our MEW fiber scaffolds are capable of containing bulk hydrogels within the femoral bone defect yet are highly porous to facilitate cell and tissue infiltration, potentially providing further advantages in enhancing early bone repair and long-term bone remodeling. Future studies will investigate the long-term degradation profile of MEW PCL fiber scaffolds *in vivo*, while tuning the use of BMP-2 affibodies to further optimize BMP-2 release profiles and bone regeneration.

## 5. Conclusions

In this work, we engineered a MEW fiber scaffold-reinforced affibody-conjugated hydrogel to tune BMP-2 delivery and enable easy surgical implantation. Scaffold-reinforced hydrogels maintained their mechanical properties and ability to control BMP-2 through after lyophilization and rehydration, thereby demonstrating potential for further investigation into shelf stability in the future. Although conjugation of high-affinity affibodies to PEG-mal hydrogels enhanced BMP-2 retention in subcutaneous implants, affibody conjugation did not improve bone repair in rat femoral bone defects. Regardless, MEW reinforcement significantly increased bone volume and defect bridging in this model. Additional future studies will include investigating lower BMP-2 doses and a range of affibody concentrations to determine the optimal conditions to maximize local BMP-2 retention and subsequent bone regeneration within the defect. Altogether, scaffold-reinforced affibody-conjugated hydrogels show promise for locally delivering proteins in dynamic biological environments, while providing surgeons with a mechanically robust product that can be easily handled for surgery. This work is an invaluable step towards the development of hydrogel protein delivery vehicles for bone repair.

## Acknowledgements

We are grateful for funding from the National Institutes of Health (R35 Maximizing Investigators’ Research Award (MIRA), R35-GM147507) and Department of Defense (Discovery Award, Peer-Reviewed Medical Research Program W81XWH2210700), as well as support from the Lary Simpson Professorship to M.H.H. J.D. was supported by a doctoral-level post-graduate scholarship (PGS-D) from the Natural Sciences and Engineering Research Council (NSERC) of Canada. Y.C.P. was supported by the National Science Foundation Graduate Research Fellowship Program (NSF GRFP). P.C.H. was funded by the Wu Tsai Human Performance Alliance and the Joe and Clara Tsai Foundation. P.M.J. and M.A.B. were supported by the Knight Campus Undergraduate Scholars Program, University of Oregon Summer Program for Undergraduate Research, and Vice President for Research and Innovation Undergraduate Fellowship. P.D.D. was supported by the Bradshaw and Holzapfel Research Professor in Transformational Science and Mathematics. We thank members of the Hettiaratchi Lab for their thoughtful review of this manuscript and Malvika Singhal, Julia Harrer, David Frey Rubio, Juan Garcia, and Caroline Foskett for their assistance with animal surgeries. We also thank Dr. Danielle Benoit for the use of her lab’s peptide synthesizer and HPLC system.

