## Supplemental Information for "Melt Electrowritten Scaffold-Reinforced Affibody-Conjugated Hydrogels for Controlled Bone Morphogenetic Protein-2 Delivery"

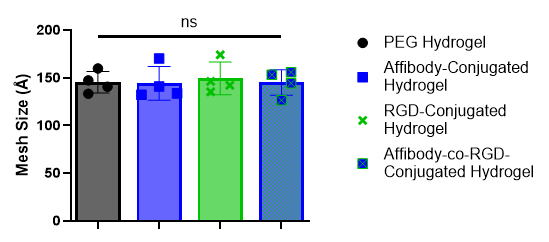

**Figure S1: Conjugation of affibodies and RGD peptides to maleimides on PEG-mal hydrogels does not affect hydrogel mesh size.** The Equilibrium Swelling Theory was used to determine the effect of affibody and RGD peptide conjugation on hydrogel mesh size. Mesh sizes were comparable between hydrogels and similar to other reported mesh sizes for 4-arm 20 kDa PEG-mal hydrogels. Statistical significance between groups was determined using one-way ANOVA with Tukey’s post-hoc test. n=4. ns= not significant.

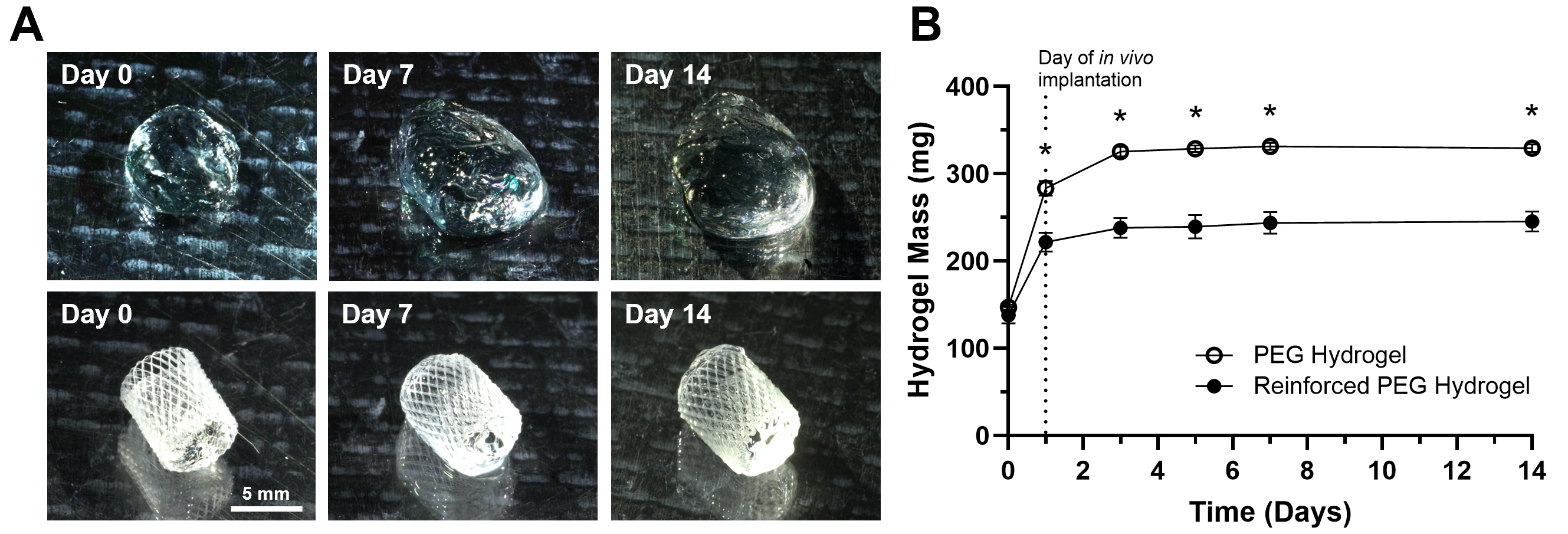

**Figure S2: Effect of MEW scaffold reinforcement on swelling and stability of PEG-mal hydrogels over 14 days.** Hydrogels with and without MEW scaffold reinforcement were initially swollen overnight (Day 0-1) in 0.1% BSA in PBS and then incubated at 37 °C and weighed and imaged periodically over 14 days. A) Images of hydrogels with and without MEW scaffold reinforcement before swelling (Day 0), at Day 7, and at Day 14. Scale bar = 5 mm. B) Mass of non-reinforced and reinforced PEG-mal hydrogels over 14 days. Hydrogels were implanted *in vivo* at Day 1 after swelling overnight. Statistical significance between groups was determined by two-way ANOVA with Tukey’s post-hoc test. n=3. * p<0.05.

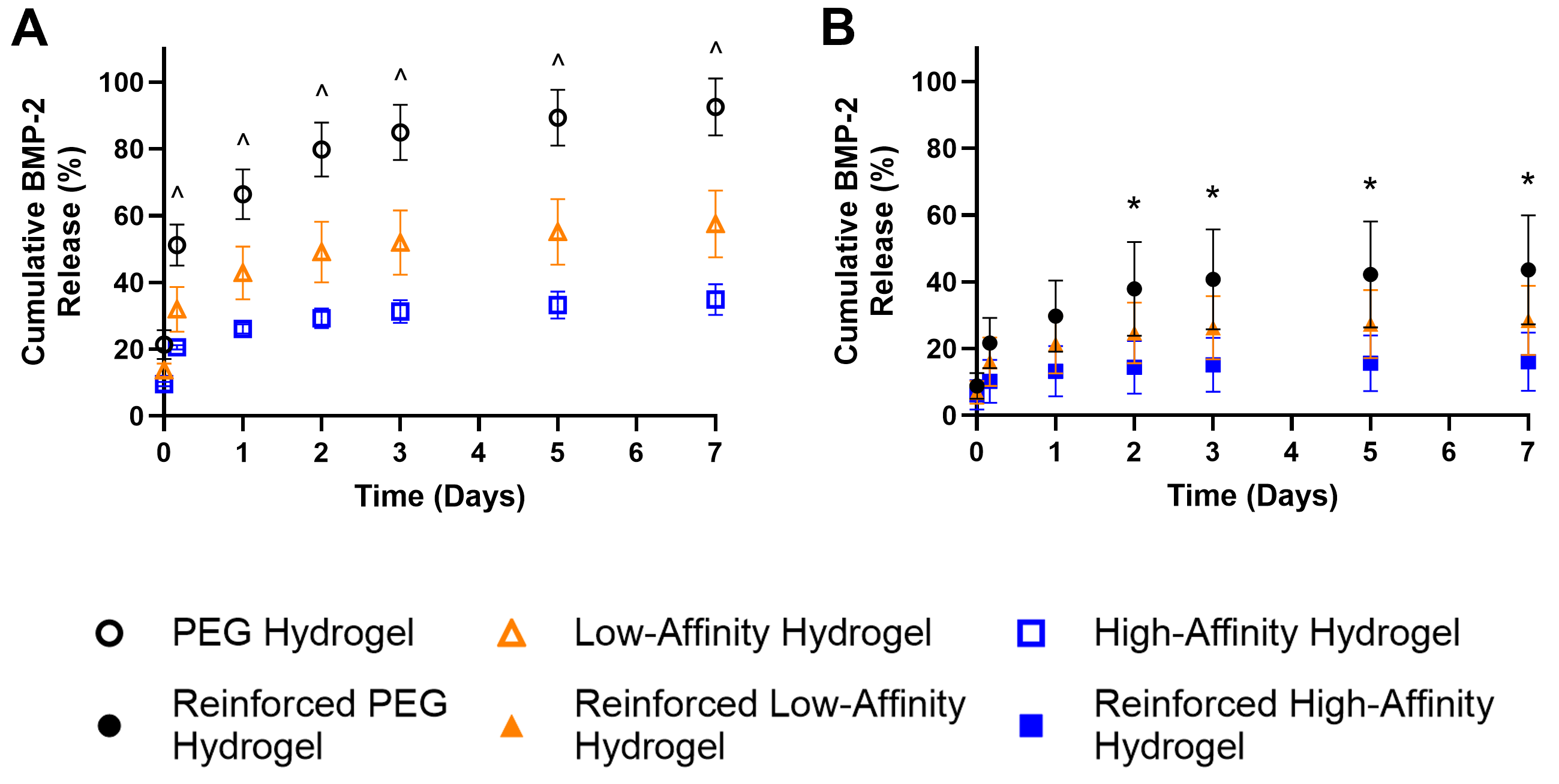

**Figure S3: Effect of BMP-2-specific affibodies on BMP-2 release.** 100 ng of BMP-2 were loaded into PEG-mal hydrogels with or without MEW scaffold reinforcement and with or without conjugated low- or high-affinity BMP-2 affibodies. BMP-2 release into 10% (v/v) FBS in PBS was monitored over 7 days. A) Release of BMP-2 from hydrogels without MEW scaffolds over 7 days. B) Release of BMP-2 from hydrogels reinforced with MEW scaffolds over 7 days. Statistical significance between groups at each time point was determined using two-way ANOVA with Tukey’s post-hoc test. n=3-4. * p<0.05 between PEG hydrogel vs. high-affinity hydrogel, ^ p<0.05 between all groups.

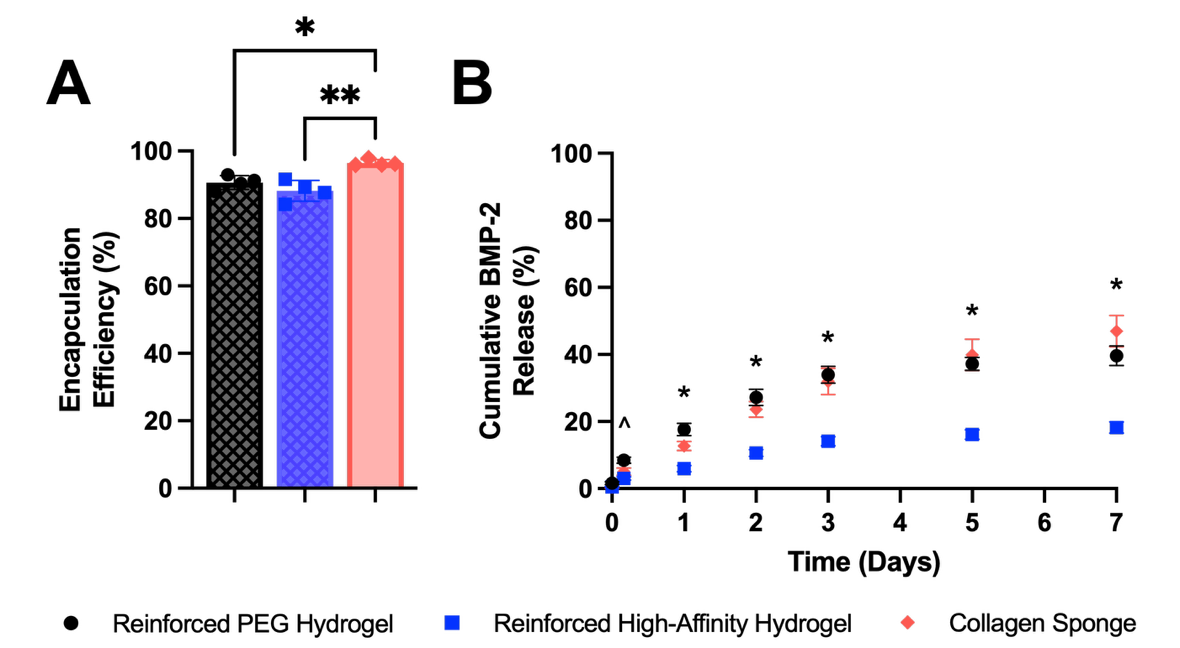

**Figure S4: Comparison of BMP-2 encapsulation and release between reinforced PEG-mal hydrogels and collagen sponges.** 100 ng of BMP-2 were loaded into collagen sponges or reinforced PEG-mal hydrogels with or without conjugated high-affinity BMP-2 affibodies. A) Initial BMP-2 encapsulation efficiency in reinforced hydrogels, reinforced high-affinity hydrogels, and collagen sponges. Statistical significance between groups was determined using one-way ANOVA with Tukey’s post-hoc test. n=4. * p<0.05, ** p<0.01. B) Release of BMP-2 from reinforced hydrogels, reinforced high-affinity hydrogels, and collagen sponges over 7 days. Statistical significance between groups at each time point was determined using two-way ANOVA with Tukey’s post-hoc test. n=4. ^ p < 0.05 between reinforced PEG hydrogel and reinforced high-affinity hydrogel, * p<0.05 between reinforced PEG hydrogel or collagen sponge and reinforced high-affinity hydrogel.

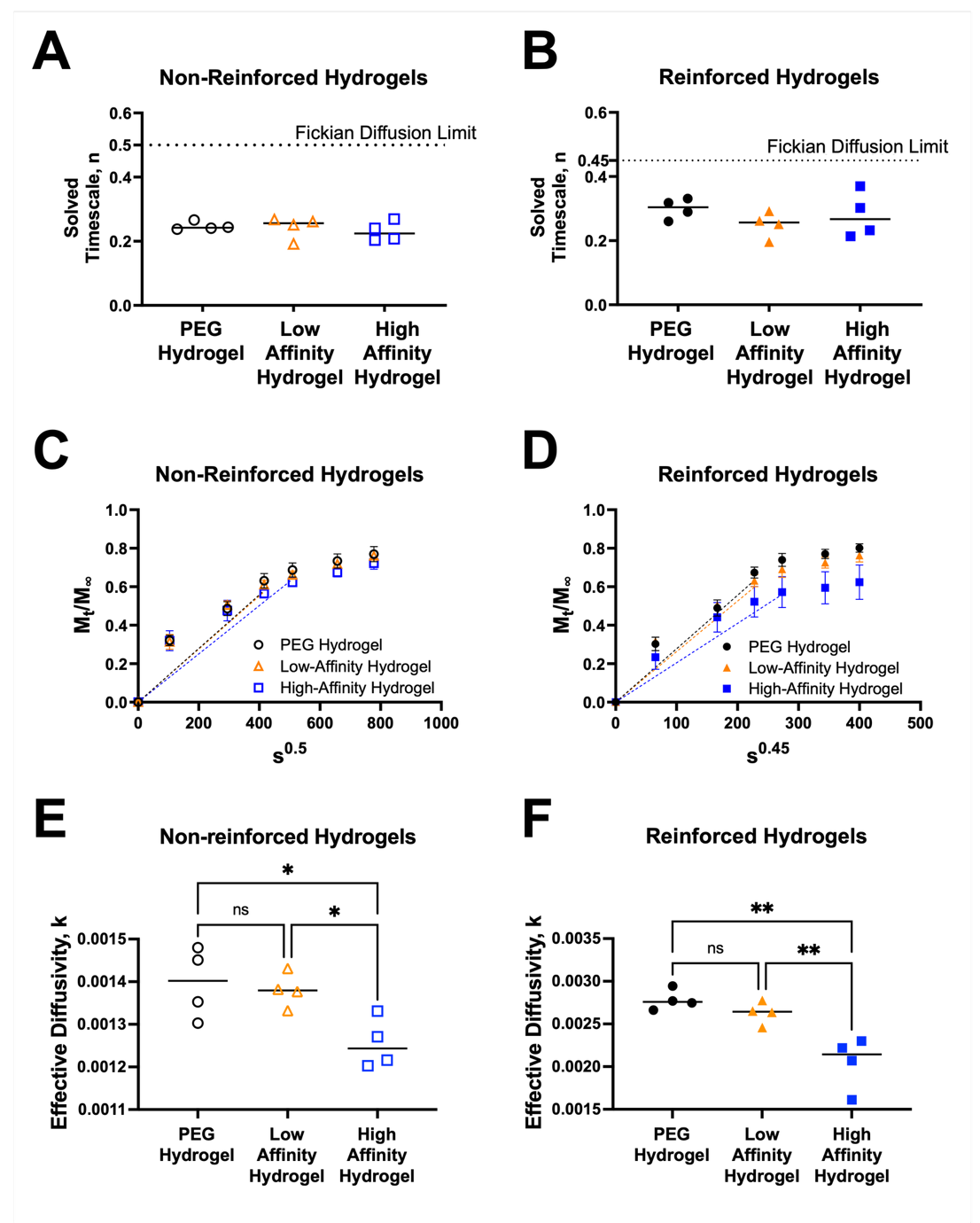

**Figure S5: Korsmeyer-Peppas model for BMP-2 release from non-reinforced and MEW scaffold-reinforced hydrogels.** Cumulative release of BMP-2 from non-reinforced and MEW scaffold-reinforced hydrogels was normalized to final cumulative release, and the data were fit to Equation 3. Calculated timescale exponent, n, for A) non-reinforced hydrogels and B) scaffold-reinforced hydrogels without affibodies, with low-affinity affibodies, or with high-affinity affibodies. The dotted lines indicate the upper limit of Fickian diffusion regimes for slab (non-reinforced) and cylindrical (scaffold-reinforced) geometries. All hydrogels exhibited Fickian diffusion. C) Normalized release of BMP-2 from non-reinforced hydrogels at timescale of s^0.5^, which is the upper limit of Fickian diffusion for a slab. D) Normalized release of BMP-2 from reinforced hydrogels at timescale of s^0.45^, which is upper limit of Fickian diffusion for a cylinder. E, F) Calculated rate of release from hydrogels with fixed timescale exponent. E) Effective diffusivity of BMP-2 from non-reinforced hydrogels with fixed timescale exponent of 0.5 for release from a slab. F) Effective diffusivity of BMP-2 from scaffold-reinforced hydrogels with fixed timescale exponent n=0.45 for release from a cylinder. Statistical significance between groups was determined using one-way ANOVA with Tukey’s post-hoc test. n=4. ns = not significant, * p<0.05, ** p<0.01.
